# Novel plasma exosome biomarkers for prostate cancer progression in co-morbid metabolic disease

**DOI:** 10.1101/2022.02.01.478722

**Authors:** Naser Jafari, Andrew Chen, Manohar Kolla, Isabella R. Pompa, Yuhan Qiu, Rebecca Yu, Pablo Llevenes, Christina S. Ennis, Joakin Mori, Kiana Mahdaviani, Meredith Halpin, Gretchen A. Gignac, Christopher M. Heaphy, Stefano Monti, Gerald V. Denis

## Abstract

Comorbid Type 2 diabetes (T2D), a metabolic complication of obesity, associates with worse cancer outcomes for prostate, breast, head and neck, colorectal and several other solid tumors. However, the molecular mechanisms remain poorly understood. Emerging evidence shows that exosomes carry miRNAs in blood that encode the metabolic status of originating tissues and deliver their cargo to target tissues to modulate expression of critical genes. Exosomal communication potentially connects abnormal metabolism to cancer progression. Here, we hypothesized that T2D plasma exosomes induce epithelial-mesenchymal transition (EMT) and immune checkpoints in prostate cancer cells. We demonstrate that plasma exosomes from subjects with T2D induce EMT features in prostate cancer cells and upregulate the checkpoint genes *CD274* and *CD155*. We demonstrate that specific exosomal miRNAs that are differentially abundant in plasma of T2D adults compared to nondiabetic controls (miR374a-5p, miR-93-5p and let-7b-3p) are delivered to cancer cells, thereby regulating critical target genes. We build on our previous reports showing BRD4 controls migration and dissemination of castration-resistant prostate cancer, and transcription of key EMT genes, to show that T2D exosomes require BRD4 to drive EMT and immune ligand expression. We validate our findings with gene set enrichment analysis of human prostate tumor tissue in TGCA genomic data. These results suggest novel, non-invasive approaches to evaluate and potentially block progression of prostate and other cancers in patients with comorbid T2D.

## 1. Introduction

The incidence of obesity-driven diabetes continues to increase worldwide, and parallel to this trend, an increase in incidence of several obesity-related cancers has been reported [1]. Already in the U.S., for example, >100 million adults have been diagnosed with diabetes or pre-diabetes [2], about 90% as Type 2 diabetes (T2D) [3], a frequent metabolic complication of obesity, emphasizing the scale of the public health challenge. The severity of comorbid T2D also predicts worse cancer outcomes; associations are well established for breast cancer [4], head and neck cancer [5], colorectal cancer [6] and several other solid tumors. In prostate [7, 8] and other cancers [9], increased burden of comorbidities also correlates with advanced age [10] and worse outcomes. Yet, the cellular and molecular mechanisms that explain these associations remain poorly understood. Although comorbidity is considered in clinical decision making, most cancer clinical trials exclude such patients, leaving us with a weak understanding of how tumor microenvironment factors may drive tumor progression. These knowledge gaps are important for medically underserved patient populations, where obesity and T2D are prevalent [11, 12], often in association with food deserts and an obesogenic built environment [13]. There is urgent need for novel, robust biomarkers to evaluate risks for cancer progression and assist clinical decision making for such underserved patients [14, 15].

New work in molecular endocrinology is revealing that cancer patients with comorbid chronic obesity and/or metabolic complications have adverse signaling in the adipose microenvironments of breast [16-19] or prostate cancers [20-22], compared to patients with the same type and stage of cancer, who are otherwise metabolically normal. In prostate cancer, obesity and its metabolic complications have been studied intensively to identify serological and histological data that would help detect cases at high risk for failure of androgen deprivation therapy (ADT), progression and metastasis in patients with these co-morbid conditions. Metabolic biomarkers that include elevated insulin-like growth factor-1, insulin and C-peptide [23, 24], leptin, glucose [25, 26], pro-inflammatory cytokines, lipid profiles [27] and androgen levels have all been associated with worse outcomes [28, 29], but opinion has long been divided about which factors are most critical. Metabolic status is also a concern because ADT is known to induce insulin resistance [30]. However, clinical trials have not yet established which serological markers of metabolism are most informative for the wide range of patients with prostate cancer on various ADT and metabolic medications, and at which stages of disease progression specific markers would have greatest value. Innovative directions would be helpful.

We took as a starting point the clinical observation that poor control of T2D in prostate cancer associates with rapid emergence of resistance to the anti-androgens abiraterone acetate and enzalutamide, compared to patients with prostate cancer who have well controlled blood glucose [26]. Non-invasive biomarkers have been explored for diagnostic, prognostic and therapeutic utility, including most recently circulating tumor DNA [31] and microRNAs (miRNA) [32]. In particular, miRNA biomarkers have gained attention because, unlike other nucleic acid biomarkers, these factors may be functional in prostate cancer [33]. Upon delivery to target tissues, miRNAs are capable of reprogramming cell metabolism and fate to affect the course of progression, metastasis and therapeutic responses. Deeper understanding of miRNA mechanism and gene targets may suggest novel therapeutics or prognostic biomarkers [34] to understand progression risks. We wondered whether circulating miRNAs might differ between T2D and non-diabetic (ND) subjects.

Significant evidence implicates exosomes as carriers of miRNAs in blood [35], saliva [36] or other body fluids that can be sampled non-invasively for biomarker assessment in cancer. Several studies have investigated blood miRNAs derived from tumors as an approach to evaluate cancer diagnosis [37, 38], prognosis [39] and recurrence [40]. However, we took a converse approach, and investigated plasma exosomal miRNAs as biomarkers of co-morbid T2D that instead might have functional impact on prostate cancer progression. Our rationale is that obesity-driven metabolic disease has long been studied in prostate cancer incidence [41, 42], progression [43] and prostate cancer-specific mortality [44], although diabetes has been shown to be protective in some prostate cancer studies [45, 46]. Despite intensive investigation, the mechanisms and clinical variables most strongly associated with incidence, progression, distant metastasis and mortality [41, 47], and the impact of metabolic medications [48], have not been settled. We considered that plasma exosomes in subjects with obesity-driven T2D might be leveraged to assess risk of progression and treatment resistance in prostate cancer, and might have functional significance to understand mechanisms of tumor progression.

We recently showed that the metabolic status of mature adipocytes determines the payload of released exosomes. Adipocytes that have been rendered insulin resistant by exposure to pro-inflammatory cytokines, or that were isolated from adipose tissue of adult subjects with T2D, release exosomes that drive increased migration, invasiveness, epithelial-to-mesenchymal transition (EMT), gene expression associated with cancer stem-like cell formation and aggressive, pro-metastatic behavior in breast cancer cell models, compared to adipocytes that are insulin sensitive or isolated from adipose tissue of ND subjects [49]. We built on these observations and hypothesized that similar phenotypes would be observed for plasma exosomes from T2D subjects compared to ND controls, using prostate and breast cancer cell lines as a readout.

Here, we report that exosomes from peripheral blood plasma of subjects with T2D induce EMT in DU145 cells, a model for hormone-refractory and aggressive prostate cancer. Additionally, we tested the upregulated T2D exosomal miRNAs on *SNAI1* and *AHNAK* expression in PC3 and DU145 prostate cancer cells, as representative EMT genes. We performed immunofluorescence imaging to verify vimentin expression as a protein marker of EMT in these two cell types, and verified results in 22Rv1 and LNCaP systems as additional forms of androgen receptor-expressing prostate cancer models.

Surprisingly, exosomes purified from the plasma of ND subjects suppressed transcription of classical EMT genes in the same model, suggesting that ND exosomes may harbor cancer chemopreventive miRNAs. We noted in the same system that T2D exosomes also induce expression of the immune checkpoint gene *CD274*, which encodes the immune checkpoint protein PD-L1. We had previously shown that the somatic members of the Bromodomain and ExtraTerminal (BET) family of proteins (BRD2, BRD3, BRD4) are positive co-regulators of the PD-1/PD-L1 axis in triple negative breast cancer models [50]. We have also shown that BRD4 regulates migration and dissemination of castration-resistant prostate cancer and transcription of key EMT genes [51, 52]. Using the pan-BET inhibitor JQ1 and the BRD4-selective PROTAC degrader MZ-1, we further demonstrate here that BRD4 is a critical effector for plasma exosome-driven prostate cancer aggressiveness, and functionally couples EMT and immune checkpoint gene expression in prostate cancer. Functional miRNAs that block pro-metastatic behavior of tumor cells could be valuable tools for discovery of novel therapeutics for advanced cancers. Our results lay the groundwork for a deeper, clinical translational investigation of plasma exosomes as functionally critical drivers of prostate tumor progression in patients with comorbid T2D, and as potential biomarkers that are both robust and non-invasive.

## 2. Materials and Methods

### 2.1. Cell lines and reagents

Cell lines were as previously described [51, 52]. The DU145, 22Rv1 and LNCaP prostate cancer cell lines were cultured in RPMI-1640 medium (Gibco). PC3 cells were cultured in F12K (Kaighn’s) medium (Gibco, 21127022). All culture media were supplemented with 10% fetal bovine serum (FBS, Corning) and 1% antibiotics (penicillin/streptomycin, Gibco). As positive controls for induction of EMT genes or PD-L1, we treated the cells with transforming growth factor (TGF)-β or interferon (IFN)-γ (5 ng/mL), respectively, as we previously published [53]. Paraformaldehyde solution (AAJ19943K2, Thermo Scientific) and 4′,6-diamidino-2-phenylindole dihydrochloride (DAPI; FluoroPure grade; D21490, Thermo Scientific) were used to fix and stain the nuclei. Recombinant Human Interferon gamma protein (Active) (ab259377), and Recombinant human TGF beta 1 protein (ab50036) were purchased from Abcam.

### 2.2. RNA staining and immunofluorescence imaging

Exosomal RNA was stained using SYTO™ RNASelect™ from Thermofisher (cat number: S32703) according to the manufacturer’s protocol. Alexa Fluor™ 568 Phalloidin (Thermofisher, A12380) was used to stain the actin fibers. Cellular nuclei were stained by DAPI, FluoroPure™ grade (Thermofisher, D21490).

### 2.3. qRT-PCR

Procedures were as previously described [51, 52]. Briefly, total RNA was extracted from tumor cells using an RNAeasy Kit (Qiagen). Reverse transcription reactions were performed with 1 μg of total RNA with the QuantiTect Reverse Transcription kit (Qiagen). Gene expression was measured using TaqMan™ master mix (Thermofisher, 4369510) and human gene probes as follows: *SNAI1* (Hs00195591_m1), *SNAI2* (Hs00950344_m1), *CDH1* (Hs01023895_m1), *ACTB* (Hs00357333_g1), *CD274* (encodes PD-L1, Hs01125301_m1), *CD155* (encodes PVR or TIGIT ligand, Hs00197846_m1), *VIM* (Hs00958111_m1), *TGFB1* (Hs00998133_m1), *TWIST1* (Hs01675818_s1) and *AHNAK* (Hs01102463_m1). Methods and primers were as previously published [49].

### 2.4. PCR array

RNA was isolated using QuantiTect Reverse Transcription Kit (Qiagen), and 1 µg of each sample was used to prepare 20 µL cDNA using RNeasy Plus Mini Kit (Qiagen). Human RT^2^ Profiler™ PCR Array, including eighty-four Epithelial to Mesenchymal Transition (EMT) genes (PAHS-090Z), Cancer Stem Cell genes (PAHS-176ZC), and miRNA Array were purchased from Qiagen and used as previously published [49]. For human immune exhaustion genesets used in informatics analyses, we chose PD-1 (*PDCD1*), PD-L1 (*CD274*), CTLA4 (*CD152*), TIM-3 (*HAVCR2*), *TIGIT, ICOS, TNFRSF4, CD27*, B- and T-lymphocyte attenuator (*BTLA*), *ADORA2A, CD40LG* and *CD28*. Reverse transcription reactions were performed with RT^2^ SYBR Green ROX qPCR Mastermix (Qiagen). Exosomal miRNAs were profiled using Human Serum/Plasma miRCURY LNA miRNA PCR array (Qiagen, #YAHS-106Y, Plate Format: 2 × 96-well).

### 2.5. Gene expression analysis

All C_t_ values of the genes were normalized to the respective *ACTB* gene (ΔC_t_). Then ΔC_t_ of each gene was subtracted from the control gene ΔC_t_ (Δ.ΔC_t_). For the control groups Δ.ΔC_t_ was calculated using this formula: ΔΔC_t_ = ΔC_t_ (C1 or C2 or C3) – ΔC_t_ (control average). Then, fold change was calculated using 2^-ΔΔC_t_, (2 to the power of negative ΔΔC_t_).

Next, the Z score was calculated based on this formula:

Z score = (x-mean)/SD, in which X is the fold change. Bio Vinci software (San Diego, CA) was used to cluster the genes. In order to predict disease and function, data were analyzed through the use of Ingenuity Pathway Analysis, IPA (QIAGEN Inc., https://www.qiagenbioinformatics.com/products/ingenuity-pathway-analysis).

### 2.6. Exosome isolation and characterization

Patient whole blood was obtained commercially from Research Blood Components, LLC (Watertown, MA), preserved on ice packs. Table 1 shows blood donor demographic information (Table 1). Samples were centrifuged (16k rpm, 4°C, 30□) to separate plasma, which was clarified again by centrifugation and filtration (0.2 µm) to remove large vesicles or apoptotic bodies. Exosomes were purified by size exclusion using qEV columns by automatic fractionation collector. The columns were packed with spherical beads (35nm pore size) such that the spaces between the pores fractionate exosomes with size range of 35nm-150nm. The exosomes were eluted using PBS with 1mM EDTA and stored at 4°C for exosome size distribution and concentration measurements using a NanoSight NS300 system. Exosomal preparations underwent quality control analysis as previously published [49]. T2D and ND plasma origin exosomes were normalized to 10^9^ particles added per cell culture well.

**Table 1.** Demographic information of blood donors.

| **ID** | **Age** | **Gender** | **Ethnicity** |
| --- | --- | --- | --- |
| ND #1 | 31 | Male | White |
| ND #2 | 39 | Male | White |
| ND #3 | 47 | Male | White |
| ND #4 | 22 | Female | South Asian |
| ND #5 | 23 | Male | Asian |
| T2D #1 | 56 | Female | African American |
| T2D #2 | 37 | Female | African American |
| T2D #3 | 53 | Female | African American |
| T2D #4 | 64 | Female | White |
| T2D #5 | 64 | Female | African American |

### 2.7. MiRNA profiling of plasma exosomes

Exosomal RNA was isolated using exoRNeasy Midi Kit (Qiagen, 77144). For miRNA-specific first-strand cDNA synthesis, miRCURY LNA RT Kit (Qiagen, 339340) was used. After cDNAs were synthesized, the PCR array analysis was conducted using miRCURY LNA SYBR Green PCR Kit (Qiagen, 339346) and Human Serum/Plasma miRCURY LNA miRNA PCR array (Qiagen, Catalog Number: YAHS-106Y, Plate Format: 2×96-Well) to detect plasma exosomes.

### 2.8. miRNA delivery to DU145 cells

Differentially expressed miRNAs in T2D exosomes were transfected using Lipofectamine RNAiMAX Transfection Reagent (13778150, Invitrogen). The human homologs of the synthesized miRNA mimics were purchased from Thermofisher. The miRBase Accession numbers are as follows: hsa-miR-374a-5p (MIMAT0000727), hsa-miR-93-5p (MIMAT0000093), hsa-let-7b-3p (MIMAT0004482), and hsa-miR-424-5p (MIMAT0001341). After transfection, cells were incubated for 48 hr at 37°C, whereupon cellular total RNA was isolated and expression of *SNAI1, CD274* (encoding PD-L1), *CDH1* or other genes compared to *ACTB* control was analyzed in transfected cells.

### 2.9. Flow cytometry analysis

Single-cell suspensions were washed after collection and stained in ice-cold Ca^+2^/Mg^+2^-free PBS with a viability dye (Zombie NIR, BioLegend) for 20 min at 4°C in the dark. Cell suspensions were then washed twice with ice-cold flow cytometry buffer (Ca^+2^/Mg^+2^-free PBS, supplemented with 2% FBS and 2 mM EDTA). Cell suspensions were then stained for PD-L1 (PE-conjugated, BD Biosciences). All cell suspensions were washed twice in ice cold flow cytometry buffer before analysis. Unstained cells and single-stained controls were used to calculate flow cytometry compensation. Data acquisition (typically 1 million events) was performed on a BD LSRII instrument at the Boston University Flow Cytometry Core Facility. Data analysis was carried out using FlowJo Software (version 10.6.1, Tree Star).

### 2.10. RNA sequencing

Total RNA was isolated from DU145 cells or PC-3 cells that were untreated controls or were treated with exosomes isolated from the plasma of three ND subjects or three T2D subjects. Each experimental group was represented in biological triplicate. RNA sequencing workflow was conducted by Boston University School of Medicine Microarray and Sequencing Core. FASTQ files were aligned to human genome build hg38 using STAR [54], (version 2.6.0c). Ensembl-Gene-level counts for non-mitochondrial genes were generated using featureCounts (Subread package, version 1.6.2) and Ensembl annotation build 100 (uniquely aligned proper pairs, same strand). Separately, SAMtools (version 1.9) was used to count reads aligning in proper pairs at least once to either strand of the mitochondrial chromosome (chrM) or to the sense or antisense strands of Ensembl loci of gene biotype “rRNA” or of non-mitochondrial RepeatMasker loci of class “rRNA” (as defined in the RepeatMasker track retrieved from the UCSC Table Browser). FASTQ quality was assessed using FastQC (version 0.11.7), and alignment quality was assessed using RSeQC (version 3.0.0).

Variance-stabilizing transformation (VST) was accomplished using the “Variance Stabilizing Transformation” function in the DESeq2 R package (version 1.23.10) [55]. (Principal Component Analysis (PCA) was performed using the prcomp R function with variance stabilizing transformed (VST) expression values that were z-normalized (set to a mean of zero and a standard deviation of one) across all samples within each gene. Differential expression was assessed using the likelihood ratio test and Wald test implemented in the DESeq2 R package. Correction for multiple hypothesis testing was accomplished using the Benjamini-Hochberg false discovery rate (FDR). All analyses were performed using the R environment for statistical computing (version 4.0.2).

### 2.11. Immunofluorescence imaging

For immunofluorescence imaging, 100,000 cells were cultured in 27-mm glass bottom dishes (Thermo Fisher Scientific, 150682). For staining, cells were washed with PBS including MgCl_2_ and CaCl_2_ and then fixed with 4% paraformaldehyde for 10 min. Then, cell membranes were permeabilized with 0.5% Triton X-100 for 15 min. Cellular binding sites were blocked with 2% (w/v) BSA (bovine serum albumin) for 50 min. Afterwards, cells were stained with vimentin primary antibody (Cell Signaling, Cat number: 5741), (1:100 dilution, v/v) for 2 hours, washed five times, and then stained with immunofluorescence conjugated secondary antibody, Alexa Fluor Plus 488 (Thermo fisher, A32731), (1:200 dilution, v/v), Alexa Fluor™ 568 phalloidin (Thermo fisher, A12380), (1:1000 dilution, v/v), and DAPI counterstain as we previously published [49]. Finally, images were captured using a Nikon Deconvolution Wide-Field Epifluorescence System at the Boston University Cellular Imaging Core. Images were analyzed using ImageJ, with differences in means evaluated by comparison of a minimum of 10 individual representative cells for each condition. Immunofluorescence stain intensity was quantified by selecting the whole cell and comparing across samples. Images were assembled in Adobe Illustrator.

### 2.12. Migration assay

DU145 cells were cultured in Eagle’s Minimum Essential Medium (EMEM), supplemented with 10% FBS and 1% Penicillin-Streptomycin, and transfected with individual miRNAs, combination, or negative control miRNA (non-target) 45 nM for 48 hours. Cells were then switched to serum-free media for 3 hours and subsequently plated in 24-well, 8-µm pore size transwell plates (Thermo Fisher Scientific) for 16 hours. The cells were plated in the upper well of the transwell inserts with serum-free media, and the bottom well was filled with EMEM complete media to serve as a chemoattractant. Cells that stayed in the upper side of the membrane and did not migrate by the end of the assay were removed with a cotton swab, and the cells that migrated were fixed with ice-cold methanol for 5 min at −20°C. After fixation, cells were stained with 1% crystal violet (v/v) in 2% ethanol for 10 min at room temperature. Images were captured by an EVOS XL Core digital inverted microscope. The percentage of migration was determined by first calculating the sum of the area of total migrated/invaded cells on the entire membrane with ImageJ software (National Institutes of Health, Bethesda, MD) and then converted to relative percent migration/invasion by comparing each condition to the control condition.

### 2.13. Statistical analysis

To identify genes whose expression changes coordinately with respect to exosome treatment groups, a one-way analysis of variance (ANOVA) was performed using a likelihood ratio test to obtain a p value for each gene. Benjamini-Hochberg False Discovery Rate (FDR) correction was applied to obtain FDR-corrected p values (q values), which represent the probability that a given result is a false positive based on the overall distribution of p values. Corrected/adjusted p values such as the FDR q are the best measure of significance for a given test when many hypotheses (genes) are tested at once. The FDR q value was also recomputed after removing genes that did not pass the “independent filtering” step in the DESeq2 package. Genes with low overall expression are more strongly affected by random technical variation and more likely to produce false positive results. Wald tests were then performed for each gene between experimental groups to obtain a test statistic and p value for each gene. Statistical analyses of the *in vitro* experiments were performed using Student’s *t* test or ANOVA as indicated, and were generated by GraphPad Prism software. *p* < 0.05 was considered statistically significant.

## 3. Results

### 3.1. Exosomes from plasma of T2D subjects induce EMT in prostate cancer cell lines

We built on our previous report that exosomes from conditioned media of human mature adipocytes induce transcription of EMT genes in breast cancer cell lines. This induction is more pronounced if the adipocytes are insulin resistant or obtained from the adipose tissue of subjects with T2D [49]. First, we tested whether human plasma exosomes phenocopied this behavior, and induced EMT genes in prostate cancer cell lines. We purified exosomes from EDTA-treated, anticoagulated peripheral blood plasma of three T2D subjects and treated DU145 cells with equal numbers of exosomes (10^9^ in all cases) for two days, then isolated total RNA and assayed gene expression by RT-PCR with TaqMan probes. As hypothesized, T2D plasma exosomes induced expression of the classical EMT genes Snail (*SNAI1*) (**Fig. 1A**), *AHNAK* and vimentin (*VIM*) (**Fig. S1**), compared to equal numbers of exosomes purified from plasma of ND subjects or media (complete growth media containing RPMI-1640 + 10% FBS) control exosomes. The mesenchymal-to-epithelial (MET) gene *CDH1*, which encodes E-cadherin, behaved in the opposite manner, as expected (**Fig. 1B**). We checked the ability of T2D plasma exosomes to induce EMT genes in a different prostate cancer cell line, 22Rv1, but found that patterns were not conserved (**Fig. S2A**), which prompted deeper investigation. Having validated the DU145 cell responses by RT-PCR, we next analyzed these same total RNAs on a commercial array for well-established EMT genes [49] and observed that the T2D plasma exosomes induced a coherent, pro-EMT signature in the DU145 cells compared to ND controls (**Fig. 1C**). Of the significantly differentially expressed genes in the commercial microarray, the T2D exosomes most strongly repressed *CDH1*, in agreement with our previous report for this gene, which is associated with maintenance of the epithelial program and opposition to the mesenchymal program [49].

**Fig. 1.**
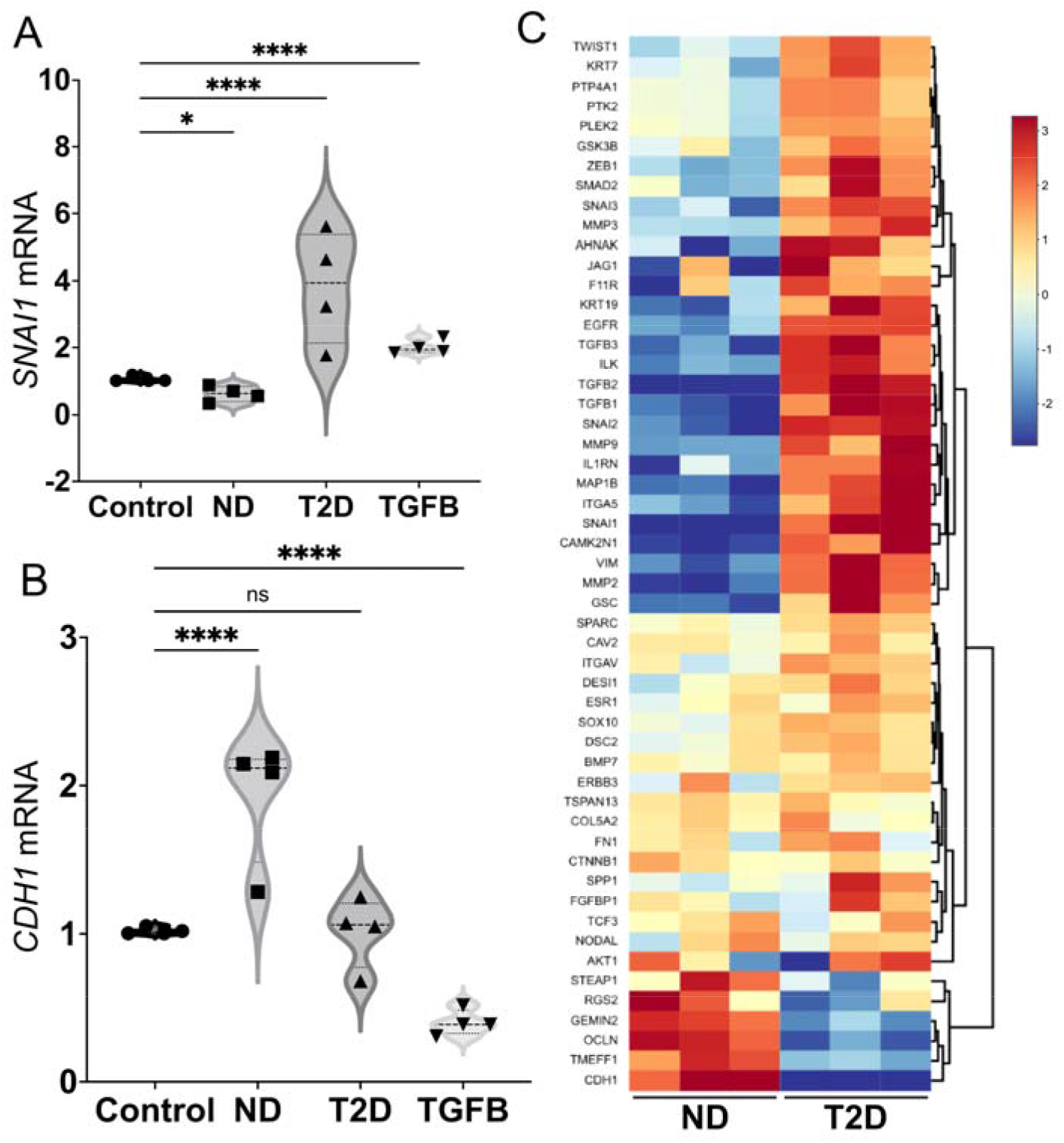
Human plasma exosomes from T2D subjects induce EMT genes in prostate cancer cell lines. The DU145 human prostate cancer cell line was treated with either ND or T2D plasma exosomes (10^9^) for 2 days. (**A**) Plasma exosomes from four, independent T2D subjects (▴) increased *SNAI1* mRNA expression after normalizing to *ACTB* of the respective sample and compared to the control cells that were treated with growth media exosomes. (•). Exosomes from four, independent ND subjects (▪) did not increase *SNAI1* mRNA expression relative to *ACTB* control. TGF-β (TGFB, 5 ng/mL) was used as a positive control (▾) for induction of pro-EMT gene transcription or repression of pro-MET gene transcription. (**B**) Plasma exosomes from T2D subjects did not induce *CDH1* expression. ND plasma exosomes increased *CDH1* mRNA expression. Data in A and B were obtained from four biological replicates of ND and T2D, and each biological replicate was conducted in three technical replicates, that are averaged in the graph. Data were analyzed by two-way ANOVA with statistical significance presented as: *, *P* = 0.0244; and ****, *P* <0.0001; *ns*, not significant. (**C**) Expression of selected EMT genes were analyzed by commercial PCR array from Qiagen. Relative expression of significantly differentially expressed genes in three independent, T2D exosome-treated samples was compared to three independent, ND exosome-treated samples. Equal numbers of exosomes (10^9^) from each sample were used. The heatmap of the PCR array result was calculated by hierarchical clustering. Scale bar (*right*) shows fold change, with red color to identify upregulation and blue color to identify downregulation. (ND, non-diabetic; T2D, Type 2 diabetic; TGFB, TGF-β positive control)

Ingenuity pathway analysis (IPA) of disease and function based on **Fig. 1C** revealed that plasma exosomes from T2D subjects strongly induced tumor cell aggressive features, such as cell spreading, protrusions, metastasis, cell motility and invasion compared to plasma exosomes from ND subjects (**Fig. 2A**), whereas pathways related to cell death by apoptosis or necrosis were downregulated by T2D exosomes, recapitulating our previous results in breast cancer models [49]. To confirm that T2D plasma exosomes also induce mesenchymal features (increased perimeter and elongation, and reduced circularity) as we previously reported [49], morphological parameters were analyzed using ImageJ. DU145 cells treated with T2D plasma exosomes showed increased perimeter and elongation, and decreased circularity, compared to cells treated with ND plasma exosomes and control cells treated with media-only exosomes (**Fig. 2BC**).

**Fig. 2.**
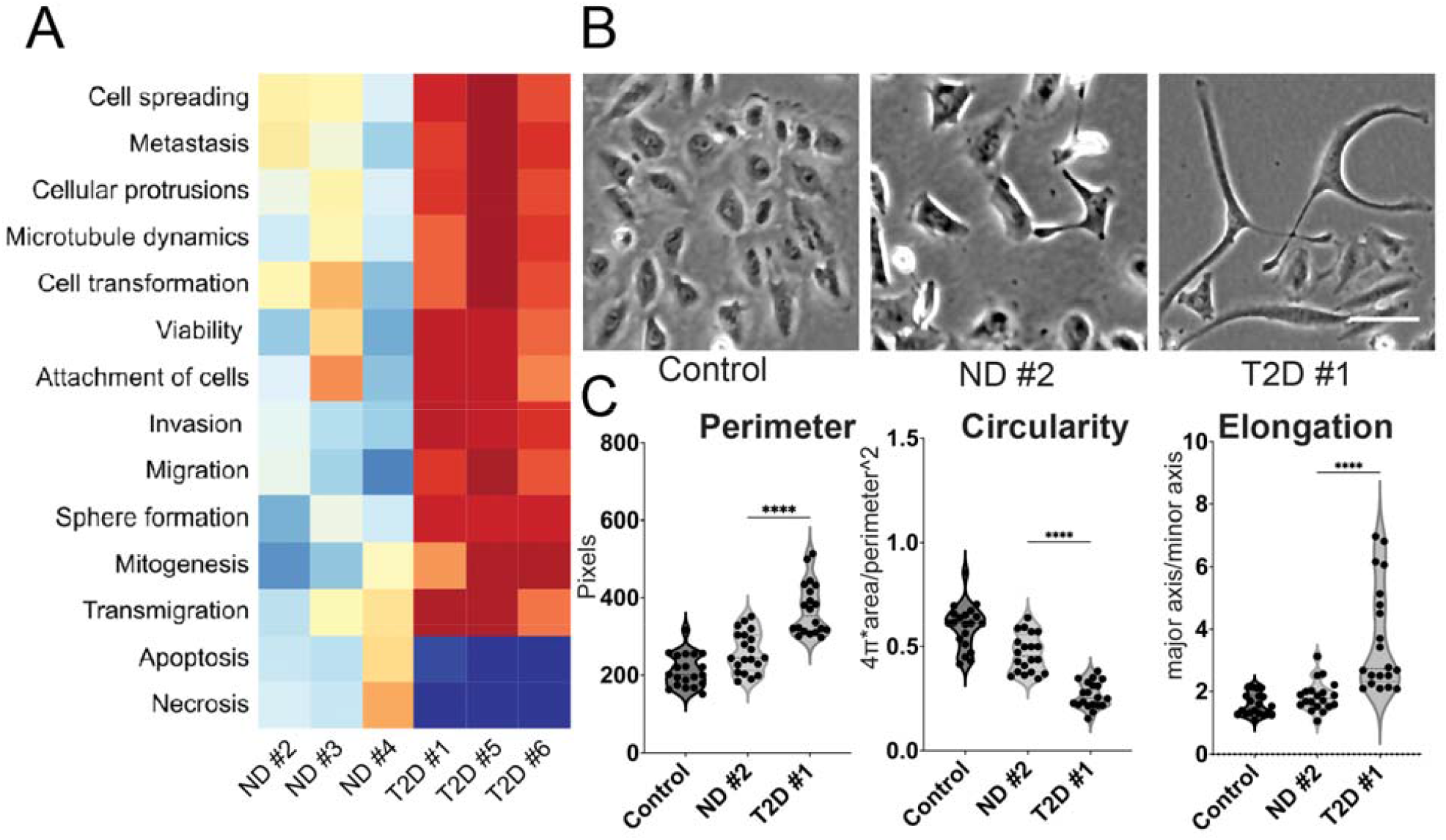
Plasma exosomes from T2D subjects induce major features of tumor cell aggressive behavior. **(A)** Ingenuity pathway analysis (IPA) of disease and function based on **Fig. 1C**. IPA prediction shows that plasma exosomes from independent T2D subjects strongly induced tumor cell signatures associated with cancer aggressiveness compared to plasma exosomes from independent ND subjects. **(B)** Morphology of DU145 cells treated with equal number of plasma exosomes ND and T2D (10^9^) compared to the control cells treated with growth media (RPMI-1640 + 10% FBS) exosomes. One representative field of view is shown, out of 25 images collected for each of the three experimental conditions with three replicates. Scale bar, 20 µm. **(C)** Quantification of cell morphology, including cellular perimeter, circularity and elongation (a parameter that is converse to circularity) measured in images from B. Expression in each experimental was compared to control (n = 25 cells each from N=3 independent experiments). Data were analyzed by one-way ANOVA, with statistical significance presented as: ****, p<0.0001. (ND, non-diabetic; T2D, Type 2 diabetic; Control, media-only exosomes)

### 3.2 Exosomes from plasma of T2D subjects induce PD-L1 expression in DU145 prostate cancer cells

Gene signatures of EMT have been associated in several tumor types with immune infiltrates that express interferon-gamma (IFN-γ)-induced genes, and correspondingly with elevated expression of immune checkpoint proteins, such as PD-L1 [56-58]. These associations prompted us to explore whether, in addition to inducing EMT networks, T2D plasma exosomes also upregulate expression of immune checkpoint genes compared to ND plasma exosomes. We found that plasma exosomes from T2D subjects did indeed upregulate genes that encode receptors important in cancer immunotherapy, including *CD274* (**Fig. 3A**), which encodes PD-L1, and *CD155* (**Fig. 3B**), which encodes the poliovirus receptor and is associated with resistance to immune checkpoint therapy in several cancer types. Next, we analyzed the effect of T2D exosomes on DU145 cells, using a commercial array focused on genes involved in inflammation and cancer immune crosstalk. The T2D plasma exosomes induced a coherent, pro-inflammatory signature in DU145 cells compared to ND plasma exosomes (**Fig. 3C**). IPA analysis of the immune/inflammation microarray data from **Fig. 3C** showed that T2D plasma exosomes strongly upregulated major pathways associated with angiogenesis [49], immune dysfunction and tumor progression, compared to plasma exosomes from ND subjects (**Fig. S3**).

**Fig. 3.**
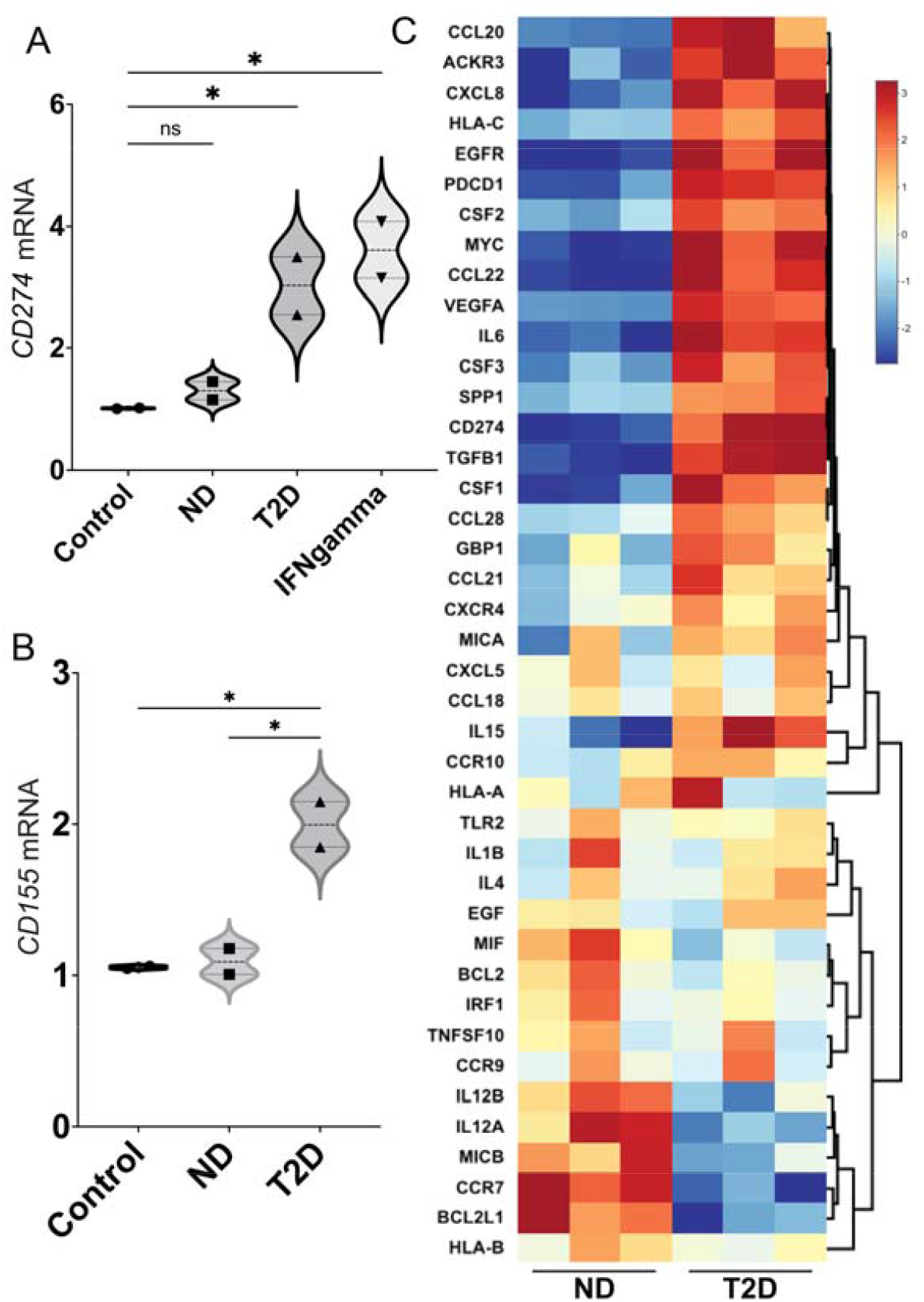
T2D plasma exosomes upregulate genes that encode immune checkpoint ligands in prostate cancer cells. Plasma exosomes from T2D subjects (▴) upregulated *CD274* (**A**) and *CD155* (**B**) mRNA expression in DU145 cells after 2 days, compared to ND control (▪). Treatment with IFN-γ (5 ng/mL) was the positive control for *CD274* induction (▾), as we previously published [53]. Treatment with growth media exosomes was the negative control (•). Data in A and B were obtained from two biological replicates of ND and T2D, and each biological replicate was conducted in three technical replicates. Data were analyzed by two-way ANOVA with statistical significance presented as: *, *P* < 0.05; *ns*, not significant. **(C)** Hierarchical clustering of genes involved in inflammation and cancer immune crosstalk, analyzed by commercial PCR array. DU145 cells were treated with plasma exosomes from three T2D subjects or three ND subjects. Equal numbers of exosomes (10^9^) from each sample were used. The heatmap of the PCR array result was calculated by hierarchical clustering. Scale bar (*right*) shows fold change, with red color to identify upregulation and blue color to identify downregulation. (ND, non-diabetic; T2D, Type 2 diabetic; Control, media-only exosomes; IFNgamma, interferon-γ)

### 3.3 MiRNA profiling of plasma exosomes

Our previous studies on payload differences between exosomes released into conditioned media by mature adipocyte cultures from T2D vs ND subjects employed mass spectrometry and proteomics to show that several proteins, particularly TSP5, were overrepresented in T2D exosomes [49]. Functional studies then showed that recombinant TSP5 alone was able to induce transcription of EMT gene signatures [49]. Therefore, we undertook a similar proteomics-based approach to investigate peptide differences between T2D and ND exosomes purified from human plasma. However, proteomics profiling of three ND and three T2D exosome isolates revealed no distinct pattern of peptides that were significantly differentially expressed between the groups (**Table S1**). We considered an alternative hypothesis that miRNAs might be differentially expressed instead of proteins, and might encode functional activities that drive the observed EMT and immune checkpoint gene expression. We compared miRNA profiles of plasma exosomes from T2D and ND subjects by a human serum/plasma miRNA PCR array and found that the metabolic differences associate with a different miRNA signature. The seven miRNAs that clustered as the most increased in T2D plasma exosomes compared to ND plasma exosomes were: miR-374a-5p, miR-93-5p, miR-28-3p, miR532-5p, miR375, miR-133b and let-7b-3p (**Fig. 4A**). Other miRNAs showed reduced differential expression in T2D plasma exosomes compared to ND plasma exosomes. The five miRNAs that clustered as the most decreased were: miR-326, miR424-5p, miR-27a-3p, miR320b and miR320d (**Fig. 4A**).

**Fig. 4.**
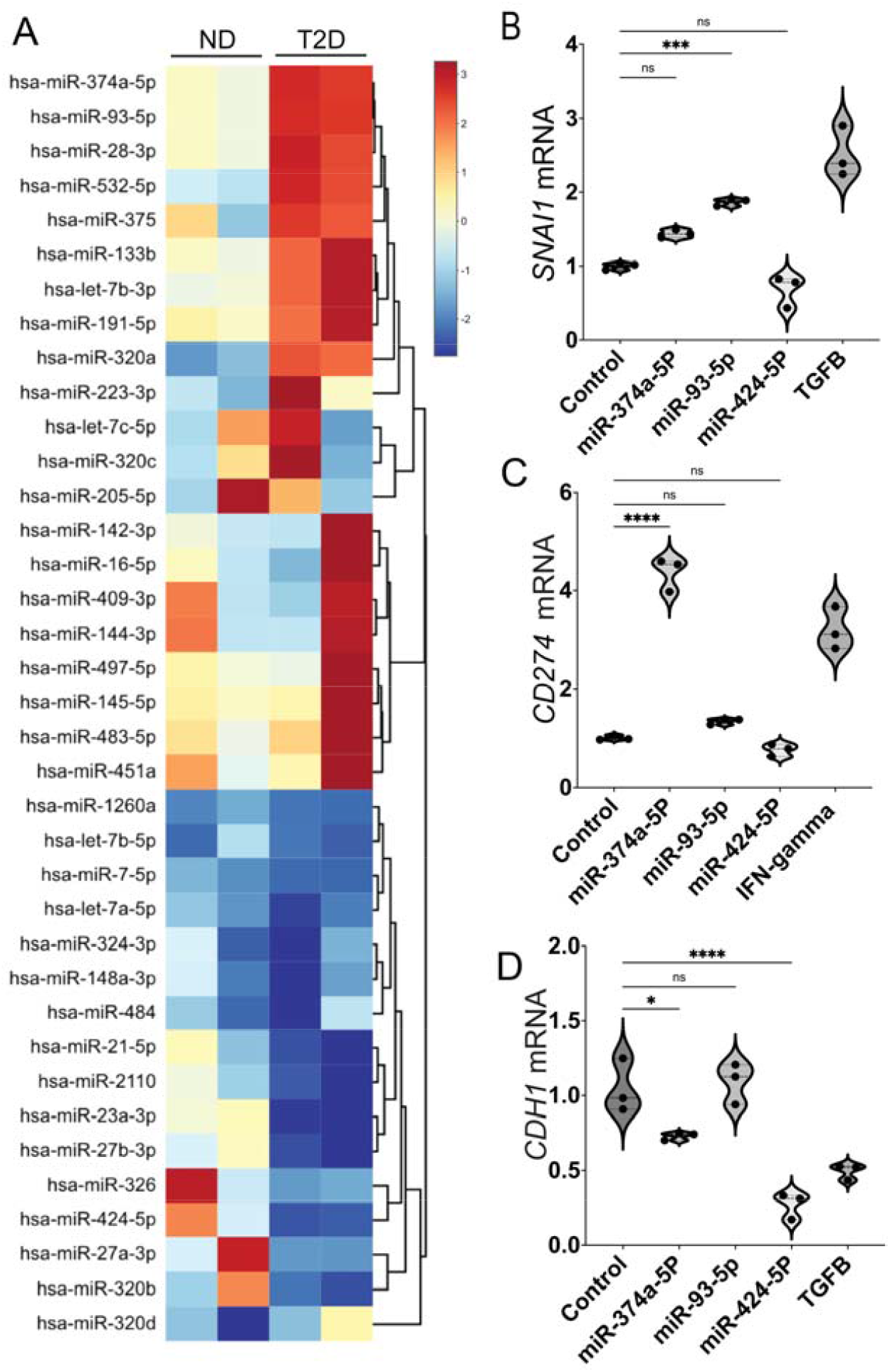
T2D plasma exosomes have distinct miRNA profile compared to ND plasma exosomes. (**A**) Heatmap of differentially expressed miRNAs in plasma exosomes of two T2D subjects compared to plasma exosomes of ND subjects. Total RNA was isolated from plasma exosomes of T2D and ND subjects, and miRNAs were profiled by a commercial PCR array. DU145 cells were transfected with individual miRNAs from (**A**) (25 nM), selected based on their functional significance for EMT and immune exhaustion. The mRNA expression of *SNAI1* (**B**), *CD274* (**C**) and *CDH1* (**D**) was tested and expression relative to β-actin (*ACTB*) is shown. Scale bar (A, *right*) shows fold change, with red color to identify upregulation and blue color to identify downregulation. Data were analyzed by one-way ANOVA with statistical significance presented as: *, *P* <0.05; ***, *P* <0.001; ****, *P* <0.0001; *ns*, not significant. (Control, media-only exosomes; TGFB, TGF-β positive control at 5 ng/mL for *SNAI1* induction; IFNgamma, interferon-γ positive control at 5 ng/mL for *CD274* induction)

For subsequent functional studies, we focused on miRNAs with known importance for cancer. Specifically, miR-374a-5p, which was highly elevated in T2D exosomes compared to ND controls, has been reported to contribute to liver cancer EMT and metastasis [59]. Additionally this miRNA is known to promote tumor progression by targeting *ARRB1* in triple negative breast cancer [60]. miR-93-5p, another miRNA that is highly expressed in T2D exosomes, has been identified as an oncogenic miRNA in a variety of human malignancies and is involved in tumor angiogenesis [61]. Bioinformatics analysis suggests that miR□93□5p has important oncogenic functions in prostate cancer [62].

A systematic reanalysis of public data revealed that let-7b is one of top five miRNAs that is consistently associated with prostate cancer progression in multiple cohorts [63]. This systematic review provides evidence from multiple publications and datasets that let-7b can serve as a biomarker for aggressive prostate cancer. Moreover, it has been shown that upregulation of let-7b in tumor-associated macrophages (TAMs) mediates angiogenesis and prostate cancer cell mobility [64]. We also examined miRNAs that were significantly under-represented in T2D exosomes compared to ND controls. Among these, miR-326 has been reported to function as a tumor suppressor in human prostatic carcinoma by targeting Mucin1 [65] and miR326 has also been shown to inhibit melanoma progression by suppressing the AKT and ERK signaling pathways [66]. These reports provided a strong rationale to test individual miRNAs for function.

We obtained three of **Fig. 4A** miRNAs from commercial sources and treated DU145 cells with 25 nM of each, for 2 days. Then total cellular RNA was isolated, cellular cDNA synthesized and fold-changes of *SNAI1, CD274* and *CDH1* were compared to *ACTB* as measured by RT-PCR with TaqMan probes (**Fig. 4BCD**). We found that miR374a-5p upregulated *CD274* (**Fig. 4C**) but not *SNAI1* (**Fig. 4B**) and slightly downregulated *CDH1* (**Fig. 4D**); miR-93-5p upregulated *SNAI1* (**Fig. 4B**) but not *CD274* (**Fig. 4C**) and did not affect *CDH1* (**Fig. 4D**); and miR-424-5p downregulated *CDH1* (**Fig. 4D**) but had no effect on *SNAI1* (**Fig. 4B**) or *CD274* **(Fig. 4C**). When tested in two other, androgen-dependent prostate cancer cell lines, 22Rv1 and LNCaP, we found similar patterns for induction of *SNAI1, TWIST1* and *CD274* (**Fig. S2BC**).

In order to partially mimic the miRNA payload of T2D exosomes, we selected upregulated miRNAs that previous publications established have a functional role in tumor cell progression. DU145 cells provide a rigorous model for migration assay under 24 hours, thus we tested the functional role of combined miRNAs on migratory potential in this model. MiR-Let-7b-3p is upregulated in T2D plasma exosomes (**Fig. 4A**), thus we tested whether miR-let-7b-3p alone plays a critical role in DU145 cell migration. A transwell assay showed that miR-let-7b-3p transfection into the cells stimulated migration (**Fig. S4**). We also tested an equimolar combination of the top three upregulated miRNAs, miR-let-7b-3p, miR-374a-5p and miR-93-5p, which were pooled and transfected into DU145 prostate cancer cells. The combination of miRNAs increased the migration of DU145 cells compared to negative control miRNA (**Fig. S4**). Surprisingly, two other upregulated miRNAs, miR-374a-5pand miR-93-5p transfected as individual miRNAs did not stimulate migration. The migration induced by the combination is attributable primarily to miR-let-7b-3p.

Next, we tested whether miR-let-7b-3p transfection also increases protein expression of vimentin, an EMT marker in DU145 cells. Immunofluorescence (IF) analysis showed that miR-let-7b-3p increased vimentin expression in DU145 cells as detected by confocal microscopy (**Fig. S5**), and although the other two miRNAs (miR-93 and miR-374) did not significantly upregulate vimentin expression, the combination of all three miRNAs at equimolar concentration still significantly upregulated vimentin. Thus, vimentin expression induced by the combination is attributable primarily to miR-let-7b-3p. We further tested whether miR-let-7b-3p upregulates transcription of selected EMT genes. qPCR TaqMan analysis showed that miR-let-7b-3p upregulated *SNAI1, AHNAK* and *CD274* (encoding PD-L1) mRNA in DU145 cells (**Fig. S6**).

To further test whether these miRNAs in combination upregulate vimentin expression in prostate cancer cells, we transfected the combined miRNAs into two prostate cancer cell lines. IF analysis showed that the three-miRNA combination increased vimentin expression in DU145 (**Fig. S5**), and PC3 cells (**Fig. S7**). Additionally, transcription assay confirmed that the combined miRNAs upregulated *SNAI1* and *Vimentin* mRNA in the same cell lines (**Fig. S8**). These results supported the overall hypothesis that T2D exosomal miRNAs are functionally active in tumor cell line models, when assayed as individual recombinant miRNAs or in combination.

We also performed a control experiment to prove that exosomal RNAs are taken up by DU145 cells. Exosome RNA was stained with RNA selective dye, then the stained exosomes were added to DU145 cells and visualized by fluorescence microscopy over several hours to track localization. Time course analysis showed that after 16 and 24 hours, the plasma exosomes RNA became concentrated in the nuclei (**Fig. S9**). Uptake reached a plateau by 16 hours.

### 3.4. RNA sequencing and Principal Component Analysis (PCA)

Next, we investigated the global transcriptional changes in prostate cancer cell lines caused by T2D exosomes in comparison to the ND exosomes. RNA sequencing and Principal Component Analysis of all signals showed that total RNA transcripts from DU145 cells treated with exosomes isolated from plasma of ND subjects had similar global transcription patterns. The transcripts from cells treated with media-only control exosomes and ND plasma exosomes clustered together well. However, transcripts from DU145 cells treated with exosomes from T2D subjects were widely separated and unique. T2D samples 1 and 2 clearly separate from the other samples along PC1 (24% of variance) and PC2 (14% of variance), respectively, suggesting that there is significant biological variability with respect to treatment with the T2D exosomes **(Fig. S10A)**. Our interpretation of this result is that ND plasma exosomes did not induce significant variation in genome-wide patterns of RNA expression compared to negative controls, but different T2D patient exosomes differ widely from ND controls, each in their own way. Furthermore, when T2D plasma exosomes are compared to ND, we identified a group of genes upregulated specifically by miR-103 **(Fig. S10B)**, that has known functional roles in EMT and immune exhaustion.

We undertook IPA analysis of differential expression of all the genes in the datasets. Network analysis revealed that miR103a and SOX2-OT induce EMT and PD-L1 expression. MiR-103a and SOX2-OT were upregulated 28.8 and 21.8 times in DU145 cells treated with T2D exosomes compared to ND exosomes (**Table S2, Fig. S10C**). Interestingly, these two genes were downregulated (5 and 3.6-fold, respectively) in the ND exosome group compared to the media-only exosome control group (**Table S2**). The RNA seq datasets were further analyzed by IPA and the potential connections of dysregulated genes and cancer cell EMT and immune exhaustion through BET proteins were explored. IPA analysis revealed that miR103a and SOX2-OT induce Cav1 and VIM respectively, which ultimately promote EMT and PD-L1 expression through BRD4 (**Fig. 5A**). Additionally, RNA seq data showed that SNAI1 and CAV1 had an upregulated trend although it was not significant (**Fig. S10B)**.

**Fig. 5.**
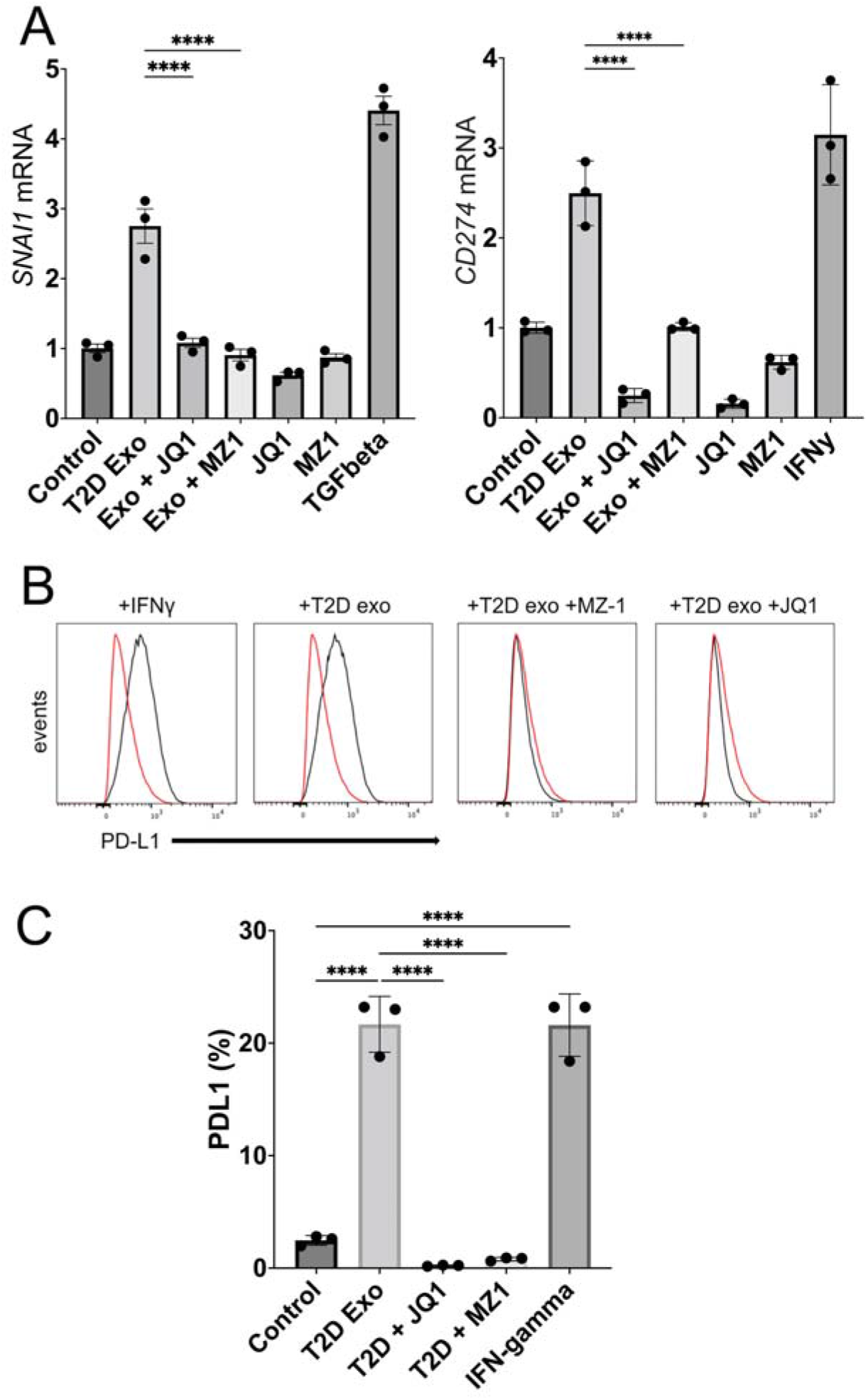
T2D plasma exosomes require BRD4 to upregulate EMT genes and PD-L1 expression. **(A)** Expression of *SNAI1* and *CD274* genes in DU145 cells measured by RT-PCR of cellular RNA after exosome treatment. T2D exosomes (10^9^; *T2D Exo*) were compared to media-only control exosomes (*Control*), or T2D Exo+JQ1 or MZ1. (JQ1 is a pan-BET inhibitor (400 nM) and MZ-1 is a BRD4-selective degrader (50 nM)). **(B)** Expression of PD-L1 was measured by flow cytometry after treatments. One million events were analyzed by Flow-Jo and presented as overlaid histograms (control, *red trace*; experimental, *black trace*). **(C)** Flow cytometry data of B were quantified with PD-L1 as percent of parent population. The experiment was conducted in triplicate with differences between means represented as bar graphs. Data were analyzed by two-way ANOVA with statistical significance presented as: ****, *P* <0.0001 (TGFB, TGF-β positive control at 5 ng/mL for *SNAI1* induction; IFNgamma, interferon-γ positive control at 5 ng/mL for *CD274* induction)

Although we showed that the top upregulated miRNAs in T2D plasma exosomes (miR-let-7b-3p, miR-374a-5p and miR-93-5p) were capable of inducing selected EMT genes, such as *SNAI1* and *VIM* alone (**Fig. 4B**; **Fig. S6**), or in combination, in both PC3 (**Fig. S8A**) and DU145 (**Fig. S8B**) cell lines, as well as Vimentin expression by IF (**Fig. S5**; **Fig. S7**) and prostate cancer cell migration (**Fig. S4**), we wanted to know whether the genome wide signature of EMT was robust across cell lines for these specific miRNAs. We therefore used RNA seq to analyze genome wide transcription of DU145 and PC3 cells alone, and as combined datasets, when challenged with the three top hit miRNAs in triplicate, compared to three non-targeted miRNAs in triplicate. Expression values for 439 differentially expressed genes in the DU145 model (**Fig. S11A**), the PC3 model (**Fig. S11B**) and both cell types combined (**Fig. S11C**) showed reproducible and coherent effects of the miRNA combination. The EMT hallmark signature was enriched in DU145 cells, PC3 cells, and when signals from both cell types were combined (**Fig. S11D**). Although the specific, representative EMT genes most strongly differentially expressed were different in each cell type, presumably because the two human prostate cancer cell lines harbor intrinsic differences, this result confirmed that the triple combination of top-hit miRNAs from T2D plasma exosomes is sufficient to induce an EMT signature in human prostate cancer cell models, interpretable by genome-wide analysis of transcription.

Our previous work has shown that the somatic BET bromodomain proteins BRD2, BRD3 and BRD4 play critical roles in transcriptional control of classical EMT genes [53, 67, 68], as well as control of immune checkpoint genes, such as *PDCD1* and *CD274*, which encode PD-1 and PD-L1, respectively. These findings have since been confirmed by others [69-71]. We therefore hypothesized here that the BET protein family functions as a central node in exosome-driven signal transduction, linking EMT to immune checkpoint function, and demonstrated here in prostate cancer cells. We have previously established in DU145 cells that low concentrations of the BRD4-selective PROTAC degrader MZ-1 (50 nM) are indeed selective to eliminate BRD4 protein while preserving BRD2 and BRD3, whereas high doses (400 nM) eliminate all three somatic BET proteins [51]. The small molecule JQ1 inhibits BET bromodomain activity through a different mechanism by competitively binding to the histone binding pocket of the bromodomain, and is not highly selective among BRD2, BRD3 and BRD4, thus we used it as a positive control for inhibition of all BET proteins, not just BRD4. We built on Fig. 4BC and measured transcription of *SNAI1* and *CD274* genes by RT-PCR. We found that MZ-1 at BRD4-selective concentrations in DU145 cells inhibited expression of *SNAI1* induced by T2D plasma exosomes, and the pan-BET inhibitor JQ1 was not able to reduce expression below the BRD4-selective level (**Fig. 5A**). These results suggested that BRD4 is required for T2D exosome induction of EMT genes. Interestingly, MZ-1 had the same inhibitory effect on CD274, but JQ1 was able to inhibit transcription to below baseline (**Fig. 5A**). We then verified by flow cytometry that PD-L1 protein expressed on the surface of treated DU145 cells was induced by T2D exosome treatment, and inhibited by MZ-1 or JQ1 (**Fig. 5BC**).

### 3.5. Gene Set Variation Analysis of TCGA prostate cancer samples

Lastly, we tested the clinical translational significance of these findings by projecting prostate cancer expression data from The Cancer Genome Atlas (TCGA) onto the EMT gene sets using Gene Set Variation Analysis (GSVA). GSVA calculates gene set enrichment for each sample independent of its labels. GSVA scores calculated with respect to the top three T2D exosomal miRNA set from above (miR-374, mi-R93, Let7b) were compared with corresponding GSVA scores calculated with respect to three EMT miRNA genesets from dbEMT2: oncogenic, tumor suppressive and dual acting (oncogenic or tumor suppressing) miRNAs in EMT. We found that GSVA scores for our T2D miRNA set were most correlated, as measured by Spearman correlation, with the oncogenic miRNAs (oncomirs) from dbEMT (coefficient of 0.5), similarly correlated with the dual acting miRNA EMT set (coefficient of 0.4), and the least correlated with the tumor suppressing miRNA EMT set (coefficient of 0.1). This result lends *in vitro* support to our overall hypothesis that the T2D exosomal miRNAs induce EMT and tumor progression.

Then, we performed Gene Set Enrichment Analysis (GSEA) on RNA seq data from DU145 cells or PC3 cells treated with the top three T2D exosomal miRNAs and observed that the EMT hallmark signature was enriched in the differentially expressed genes across both cell lines (analyzed singly or jointly) (**Fig. 6**). We also noted enrichment of gene sets associated with DNA damage, stress responses, metabolic reprogramming and angiogenesis, consistent with our previous report on the transcriptional signatures induced by T2D plasma exosomes in human breast cancer cell models [49]. Matrices to test the correlation between the signature induced by the limited miRNA show that immune exhaustion signatures were not as strongly correlated in these cell lines as the EMT gene sets (*data not shown*). These results suggest that the three miRNAs both up-regulate a similar set of genes in prostate cancer cell lines that have elsewhere been observed to be upregulated during EMT, and reflect similar EMT miRNA signatures in processes of tumor progression that are observed in TCGA clinical samples.

**Fig. 6.**
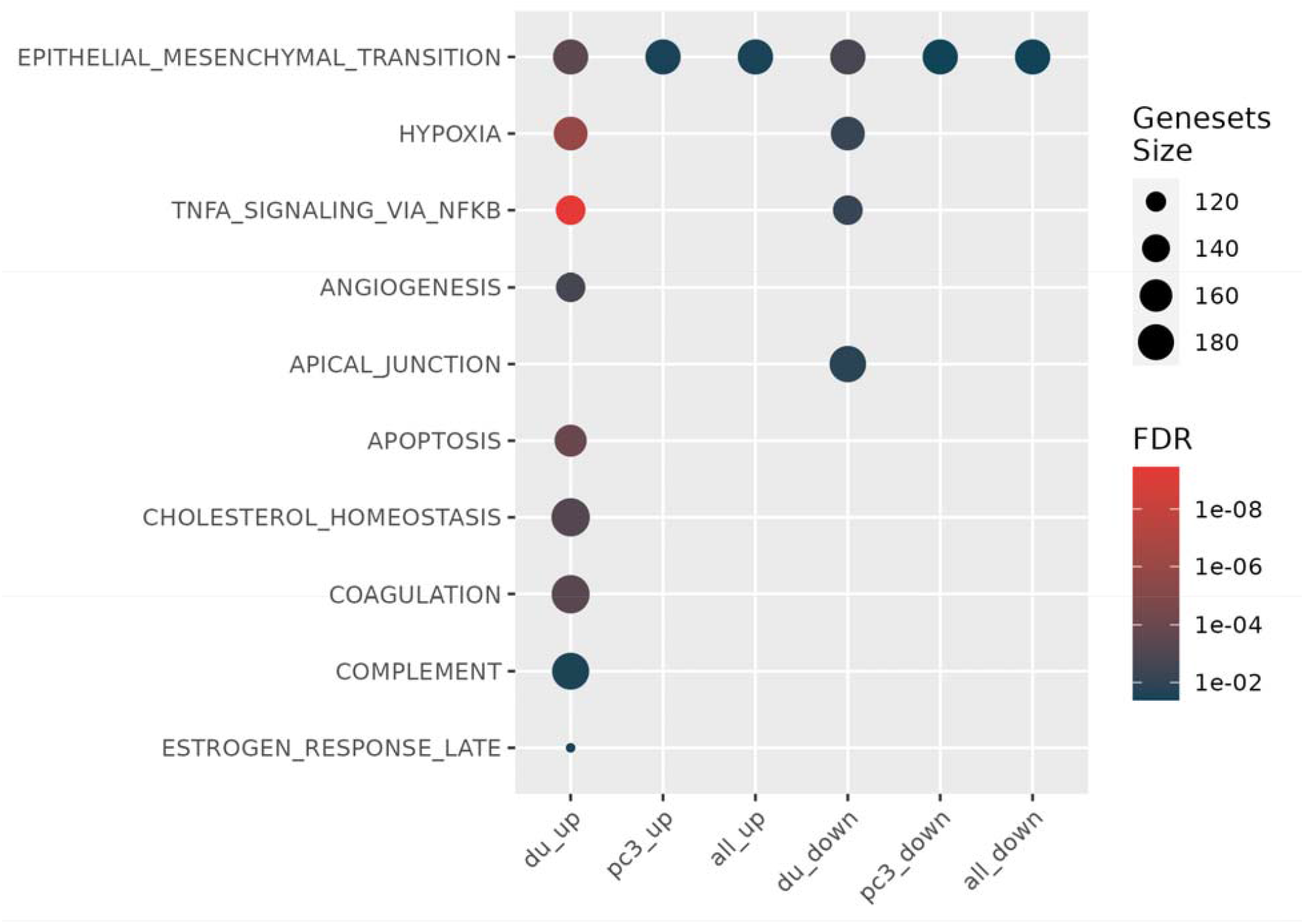
T2D exosomal miRNAs induce hallmark EMT signatures. Specific exosomal miRNAs (miR-374, miR-93, Let7b) promote tumor aggressiveness in prostate cancer cell lines, when considered as an EMT signature, shown by geneset enrichment analysis. (du_up, differentially expressed genes that showed increased expression in DU145 cells; pc3_up, differentially expressed genes that showed increased expression in PC-3 cells; all_up; the union of the DU145 and PC-3 datasets of upregulated genes; du_down, differentially expressed genes that showed decreased expression in DU145 cells; pc3<u>_</u>down, differentially expressed genes that showed decreased expression in PC-3 cells; all_down; the union of the DU145 and PC-3 datasets of downregulated genes; FDR, false discovery rate)

## 4. Discussion

Prostate cancer exhibits significant genomic and histologic heterogeneity that complicates prognostic assessment and clinical decision making. Disease can be indolent and localized, oligoclonal with non-overlapping mutational profiles among nearby clones [72], or aggressive with rapid progression and metastatic dissemination of a lethal clone [73]. A large subgroup of cases appears to be indolent at the early stage; thus, it is important to resolve the indolent cases from aggressive cases that demand immediate treatment. Although serum level of Prostate Specific Antigen (PSA) has proven utility in combination with digital rectal examination for diagnostic screening [74, 75], PSA accuracy is suboptimal to understand cancer risks [76, 77]. Additional, non-invasive biomarkers have been explored for diagnostic, prognostic and therapeutic utility, including most recently circulating tumor DNA [31] and miRNAs [32]. In particular, miRNA has gained attention because, unlike other biomarkers, these factors may be functional in prostate cancer [33], and upon delivery to target tissues, are capable of reprogramming cell metabolism, gene expression and cell fate to redirect the course of progression, metastasis and therapeutic responses. Deeper understanding of miRNA mechanism and gene targets may suggest novel therapeutics or prognostic biomarkers [34] to understand progression risks.

Here, we took a new approach of exploring plasma exosomes for functional biomarkers that could prove useful for patients with prostate cancer. This report is the first to describe how plasma exosomes from subjects with Type 2 diabetes (i) drive pro-EMT transcriptional shifts and (ii) elevate immune checkpoint expression in human prostate cancer models. Surprisingly, we also found that plasma exosomes from ND controls showed activity to downregulate certain classical, pro-EMT genes, such as *SNAI1*, and to upregulate certain classical anti-EMT genes, such as *CDH1*, in prostate cancer cell lines (**Fig. 1C**). This observation suggests that ND status may encode chemopreventive factors that are packaged into plasma exosomes that circulate and may protect against prostate cancer progression.

Our initial approach to use differential proteomics analysis of the exosomes to identify peptides with significantly different abundance between T2D and ND exosomes recapitulated our previous approach, where we used proteomic analyses of exosomes purified from conditioned media of insulin-resistant vs. insulin-sensitive mature adipocytes [49]. However, peptide differences between the two types of plasma exosomes were not significant to warrant deeper investigation. We turned instead to analysis of miRNAs to identify potential differences. Here, we found that commercial array kits were adequate to reveal strikingly different miRNA profiles.

To investigate this model, we compared the RNA seq data of DU145 cells treated with plasma exosomes from T2D and ND subjects. We found that the T2D exosomes upregulated a subset of genes that play critical roles in both EMT and immune exhaustion (**Table 2**). In order to illustrate the pathway, genes and their fold change values were imported to IPA. By using the path explorer feature of the software, the connections among upregulated genes, EMT genes and immune exhaustion genes were revealed (**Fig. 5A**). IPA output revealed that miR103 and SOX2-OT stimulate *CAV1* and *VIM* respectively. Downstream in the pathway, BRD4 acts as the critical node, and activates *SNAI1* and *CD274*, which subsequently drive EMT and immune checkpoint expression. Other studies have proven that miR-103 promotes metastasis of colorectal cancer by targeting the metastasis suppressors DAPK and KLF4 [78]. Moreover, it has been reported that tumor derived miR-103 enhances cellular motility, invasion and EMT in esophageal squamous cell carcinoma [79] and oral squamous cell carcinoma [80]. In addition to miR-103, SOX2-OT was significantly upregulated in DU145 cells treated with T2D plasma exosomes. Published studies report that SOX2-OT facilitates proliferation and migration [81], invasion and metastasis of prostate cancer cells [82]. Taken together, these data suggest a new signaling map to link the pro-EMT effect of T2D plasma exosomes with PD-L1 expression through BRD4 (**Fig. 7**).

**Fig. 7.**
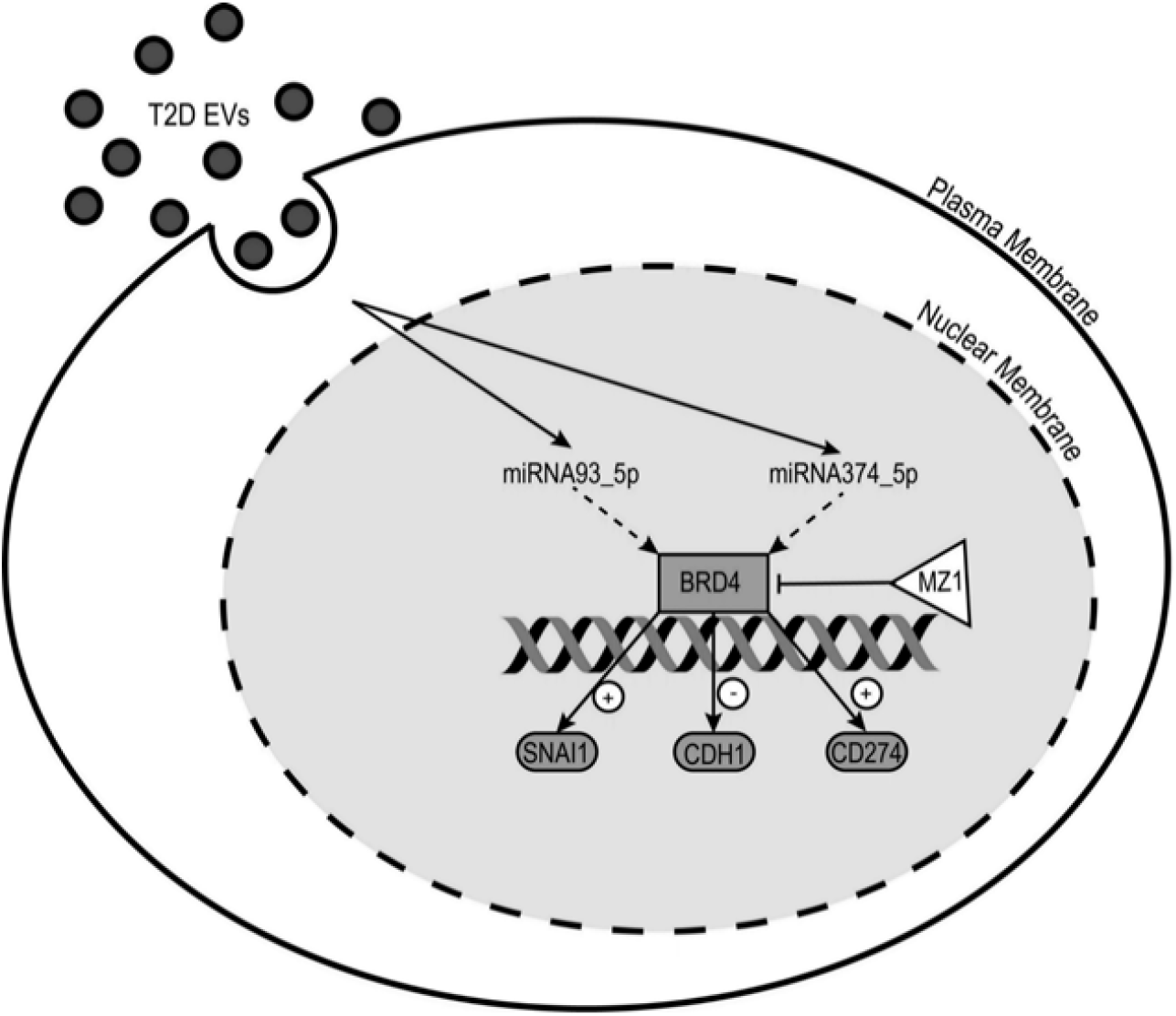
Model for T2D exosome delivery of miRNAs that reprogram transcription of tumor cell genes important for cancer aggressiveness. Circulating exosomes in plasma arrive at the tumor cell surface where they are internalized and release their cargo of miRNAs. These miRNAs are trafficked to the nucleus where they reprogram signal transduction pathways. In this case, miR-93-5p and miR374a-5p are shown signaling through BRD4, which is an essential transcriptional co-regulator of genes important for tumor cell aggressiveness, *SNAI1, CDH1* and *CD274*.

Principal Components Analysis (**Fig. S10A**) suggests that most non-diabetic patients share similar exosomal payloads and signaling to prostate cancer target cells, but patients with T2D may differ from each other, each exacerbating risk of progression in their own way. We suspect that individual-level differences among prostate cancer patients (with respect to their personalized profile of mutations, for example), and other differences in T2D status (duration of T2D since onset, degree of glucose control, patient age and other factors like metabolic medications), are likely critical contributors to personalized risk of prostate cancer progression. The deduced EMT signature of individual T2D patients likely also depends on the choice of prostate cancer cell line for the exosome readout. Sorting out these systemic differences to identify the most important factors will require a large clinical trial.

PD-L1 targeting is a major treatment strategy for several current cancer clinical trials of checkpoint inhibitors. The PD-L1 findings may have clinical relevance related to utility of checkpoint inhibitors for treatment of advanced prostate cancer. Although checkpoint inhibitors have demonstrated significant efficacy in other advanced malignancies, the outcomes in advanced prostate cancer have been disappointing. Randomized phase III studies have failed to demonstrate an overall survival benefit in advanced prostate cancer [83, 84]. These studies, as well as several smaller studies with PD-1/PD-L1 inhibitors, did demonstrate clinical benefit in a small percentage of patients, yet there is ongoing need to identify markers to predict response or resistance. As we show here, T2D may work through plasma exosome crosstalk to increase the plasticity of prostate cancer primary cells and induce immune exhaustion markers important for microenvironment interactions with tumor-infiltrating T cells. The consequences of this crosstalk for therapeutic agents, such as atezolizumab, are unknown and stratification by metabolic status in clinical trials using PD1/PD-L1 inhibitors may provide further insights.

We were also able to project TCGA data for human primary and metastatic prostate cancer onto genome wide EMT genesets of the human prostate cancer models DU145 and PC3, treated with the top three miRNAs that are upregulated in T2D plasma exosomes. We found good concordance between our induced EMT gene sets and the clinical datasets. Our hypothesis is supported: a limited set of T2D exosomal miRNAs are clinically important for prostate cancer progression and EMT. The ‘PC3 up’ and ‘DU145 up’ gene sets are similar to other established EMT gene sets. Validation of the functional importance of these miRNAs in TCGA databases for prostate cancer clinical samples confirms that our conclusions have real world importance for prostate cancer prognosis and outcomes, and are not restricted to *in vitro* or *ex vivo* model systems. Overall, we are able to make two major conclusions: 1. Specific plasma exosomal miRNAs could be useful as non-invasive, clinical biomarkers, when viewed through the lens of EMT; and 2. Specific exosomal miRNAs (miR-374, miR-93, Let-7b) promote tumor aggressiveness in prostate cancer cell lines, when viewed through the lens of EMT.

Considered together, our findings show that T2D plasma exosomes induce prominent EMT signatures and immune checkpoint markers in prostate cancer cell models. Social determinants, including the built environment, contribute to obesity, metabolic syndrome and T2D, which are known to associate with worse prostate cancer outcomes. Specifically, we have shown that poorly controlled blood glucose strongly stratifies risk of progression to resistance to the anti-androgen therapies abiraterone and enzalutamide [26]. Our results identify plasma exosomal microRNAs that are differentially expressed in blood of adults with T2D compared to non-diabetic adults, which are functional in cellular assays and drive prostate cancer aggressiveness. These features are highly correlated with progression signatures in human clinical samples. We suggest that our approach has value to identify novel blood-based biomarkers that may link obesogenic/diabetogenic built environments with biochemical progression of prostate cancer. Longitudinal monitoring of plasma exosome payloads in prostate cancer patients who are being treated with ADT may reveal important shifts in miRNA profile that could inform clinical decision making about altered risk for progression. Our study is the first report to unravel the oncogenic function of T2D plasma exosomes that could drive metastatic progression and treatment failure in castration-resistant prostate cancer in people with comorbid metabolic disease.

## Abbreviations

ADT: androgen deprivation therapy
BET: Bromodomain and ExtraTerminal
DAPI: 4′,6-diamidino-2-phenylindole dihydrochloride
EMT: epithelial-to-mesenchymal transition
FBS: fetal bovine serum
FDR: false discovery rate
IFN: interferon
IPA: Ingenuity Pathway Analysis
ND: non-diabetic
PCA: principal components analysis
PD-L1: programmed death-ligand 1
PROTAC: proteolysis-targeting chimera
PSA: prostate specific antigen
T2D: Type 2 diabetes
TGF: Transforming Growth Factor
VST: variance-stabilizing transformation

## Supplementary figures

**Supplemental Fig. S1.**
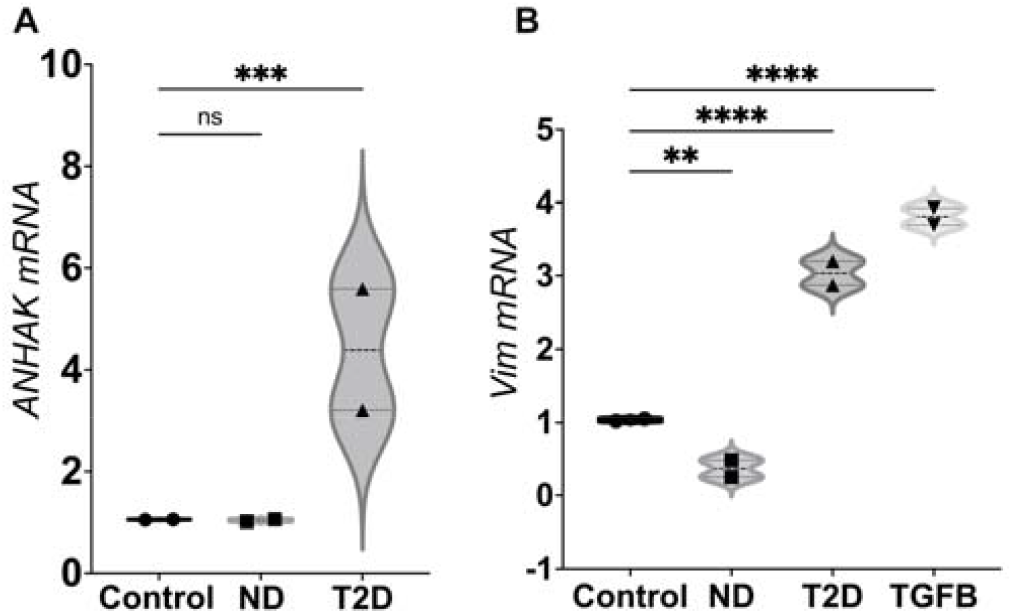
T2D exosome induction of selected EMT genes, determined by RT-PCR. Human plasma exosomes from T2D subjects (▴) induced representative EMT genes *AHNAK* (A) and *VIMENTIN* (*Vim)* (B) in DU145 cells, shown as fold-change relative to *ACTB* housekeeping gene (●). Responses for plasma exosomes from T2D subjects are compared to ND controls (▪). Data in **(A)** and **(B)** were obtained from two biological replicates of ND and T2D; each biological replicate was conducted in three technical replicates. Data were analyzed by two-way ANOVA, with statistical significance presented as: **, p< 0.01; ***, p< 0.001; ****, p<0.0001; *ns*, not significant. ND, non-diabetic; T2D, Type 2 diabetic; TGFB, TGF-β positive control (▾).

**Supplemental Fig. S2.**
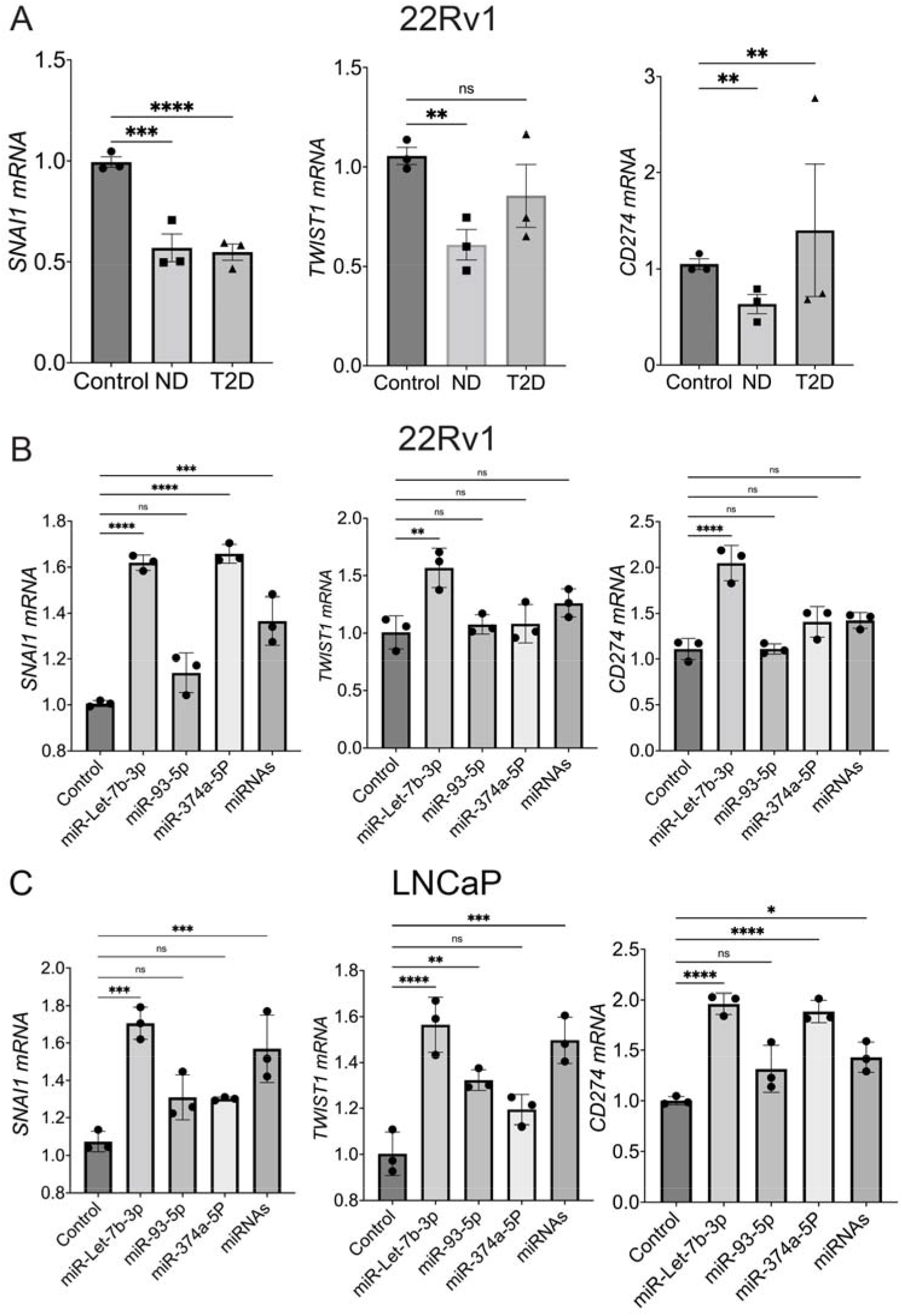
T2D exosome and exosomal miRNA induction of selected EMT and immune checkpoint genes in prostate cancer cellular models. **(A)** Effects of T2D plasma exosomes on *SNAI1, TWIST1* and *CD274* (PD-L1) in 22Rv1 cells. Cells were treated with independent exosome isolates from three T2D and three ND subjects (biological triplicate) and incubated for 48 hours. Each biological replicate was conducted in three technical replicates. **(B)** miR-let-7b upregulates selected EMT genes and *CD274* (PD-L1) in 22Rv1 cells. mRNA expression of selected genes encoding *SNAI1, TWIST1* and *CD274* was measured by RT-PCR relative to RNA encoding β-actin (*ACTB*). **(C)** miR-let-7b upregulates selected EMT genes and PD-L1 in LNCaP cells. mRNA expression of selected genes was measured by RT-PCR as in (A). (miRNAs refers to equimolar combination of three mRNAs miR-let-7b-3p, miR-374-5p and miR-93-5p (each 15 nM, total 45 nM) or negative control miRNA (non-targeting) at 25 nM. Data were analyzed by two-way ANOVA in (A), one-way ANOVA in (B), with statistical significance presented as: **, p< 0.01; ***, p< 0.001; ****, p<0.0001; ns, not significant. ND, non-diabetic; T2D, Type 2 diabetic)

**Supplemental Fig. S3.**
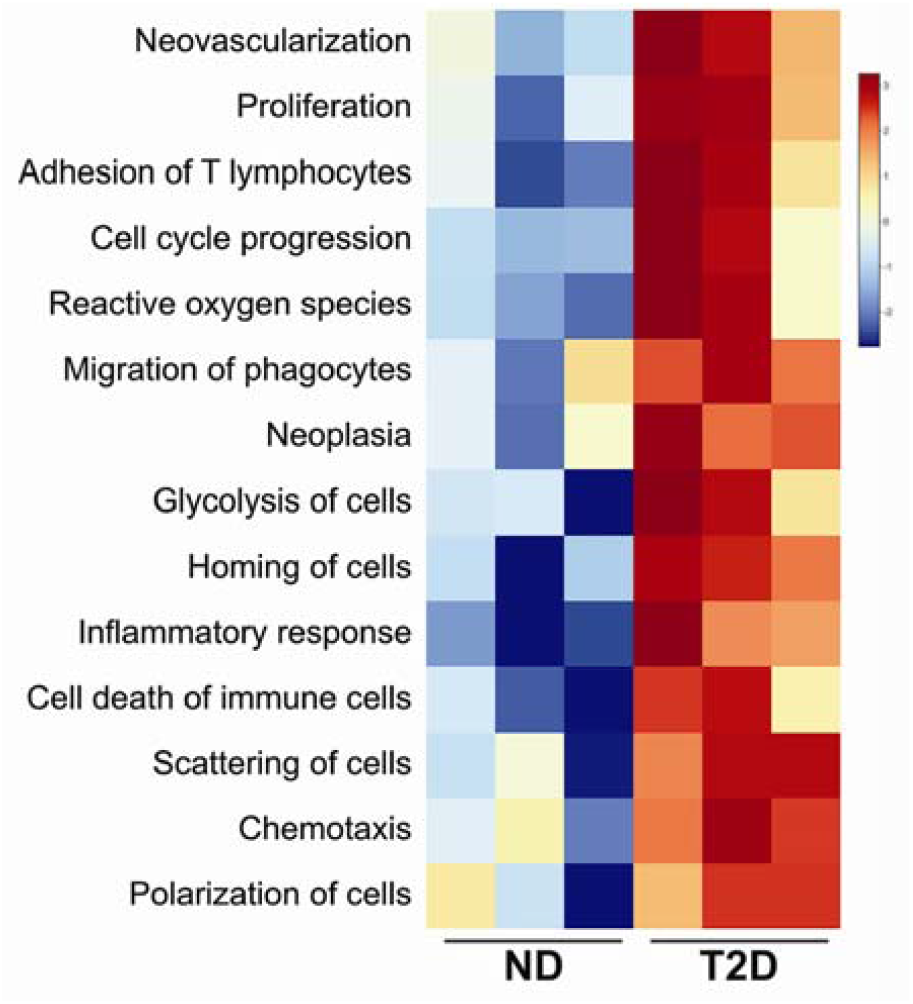
Ingenuity pathway analysis (IPA) of disease and function. IPA prediction based on Fig. 3C shows that plasma exosomes from T2D subjects induced major mechanisms associated with angiogenesis, immune dysfunction and tumor progression, compared to plasma exosomes from ND subjects. Three independent datasets for T2D are compared to three independent datasets for ND. Scale bar (*right*) shows fold change, with red color to identify upregulation and blue color to identify downregulation. (ND, non-diabetic; T2D, Type 2 diabetic)

**Supplemental Fig. S4.**
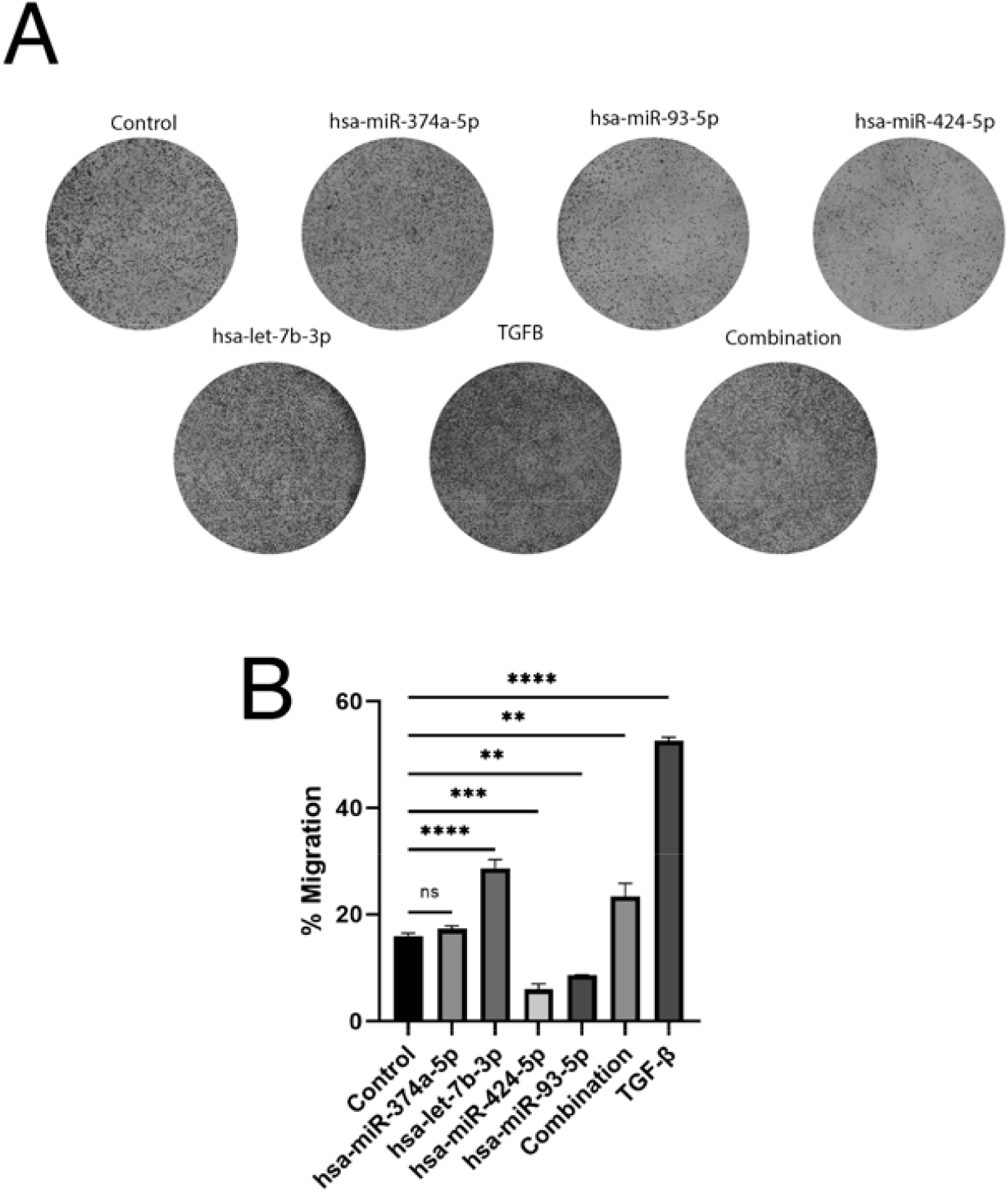
Migration assay. Migration of DU145 prostate cancer cells was measured in a 16-hour transwell assay after transfection with individual miRNAs, the triple combination, or negative control miRNA (non-target) at 45 nM each. **(A)** Cells that reached the distal side of the 8 µm pore membrane were visualized by crystal violet stain and microscopy. **(B)** Quantification of the migrated cells normalized to the background. TGFB, TGF-β positive control at 5 ng/mL for migration induction. Data are from N = 3 independent experiments. **, *P* < 0.01; ***, *P* < 0.001; and ****, *P* <0.0001 by one-way ANOVA.

**Supplemental Fig. S5.**
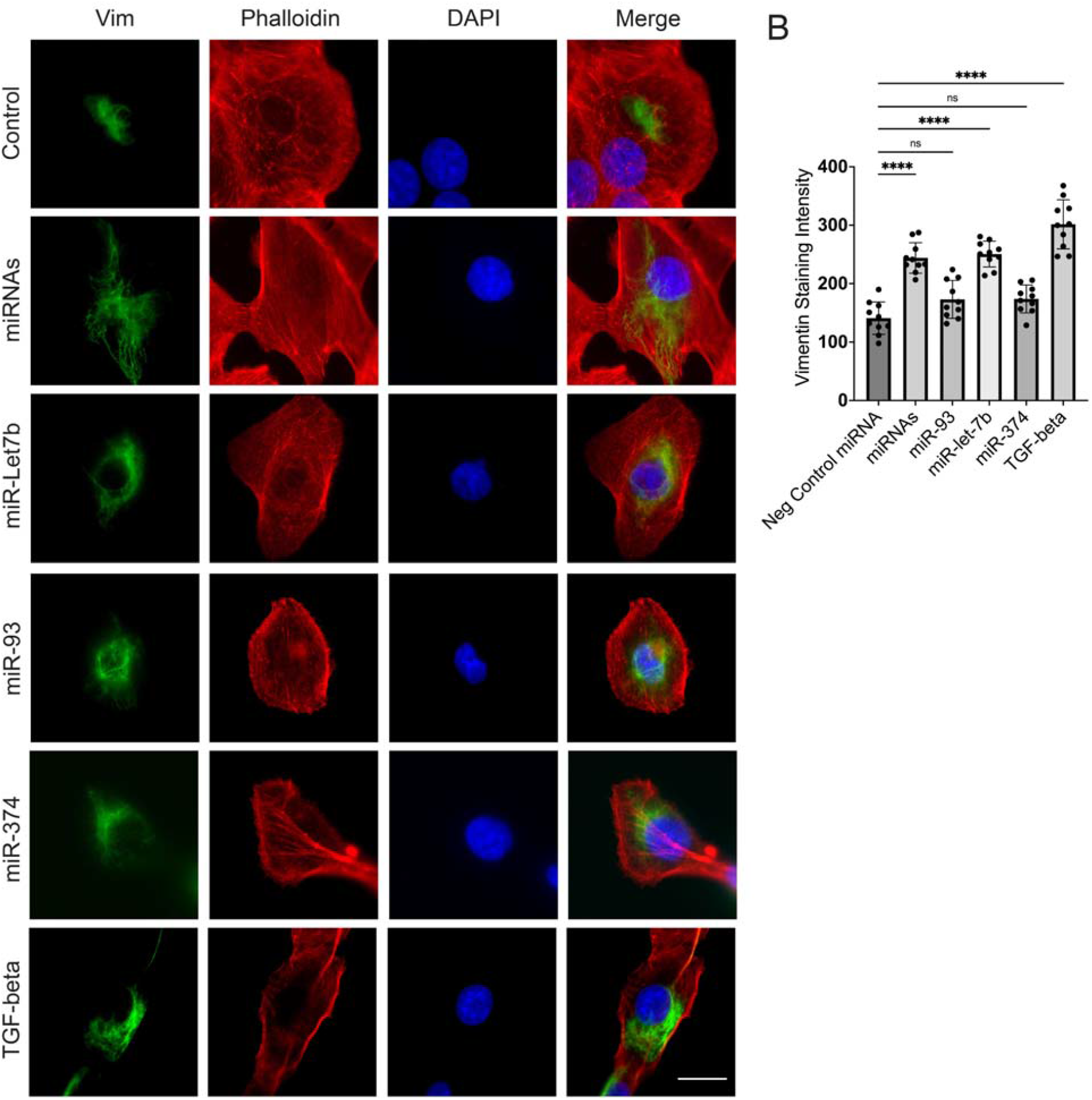
Top three exosomal miRNAs, tested individually or together for regulation of vimentin protein expression in the DU145 model. **(A)** Immunofluorescence staining for vimentin was measured in DU145 cells transfected with individual miRNAs, the combination of three miRNAs (each 15 nM, total 45 nM), or negative control miRNA (non-targeted), at 45 nM, and incubated for 48 hours. Cells were stained with vimentin primary antibody and Alexa Fluor-488 (green) secondary antibody. Cytoskeleton was stained with Alexa Fluor-568 Phalloidin (red) and nucleic DNA was stained with DAPI. Scale bar, 10 µm. For each of N = 3 independent experiments, 10 images were examined by immunofluorescence. One representative image is shown, of 10 images collected for each of the three experimental conditions with three replicates. **(B)** Quantification of vimentin staining intensity in 10 individual cells in each condition. (TGF-beta, TGF-β as a positive control. Data analyzed by one-way ANOVA, ****; *P* <0.0001, *ns*, not significant)

**Supplemental Fig. S6.**
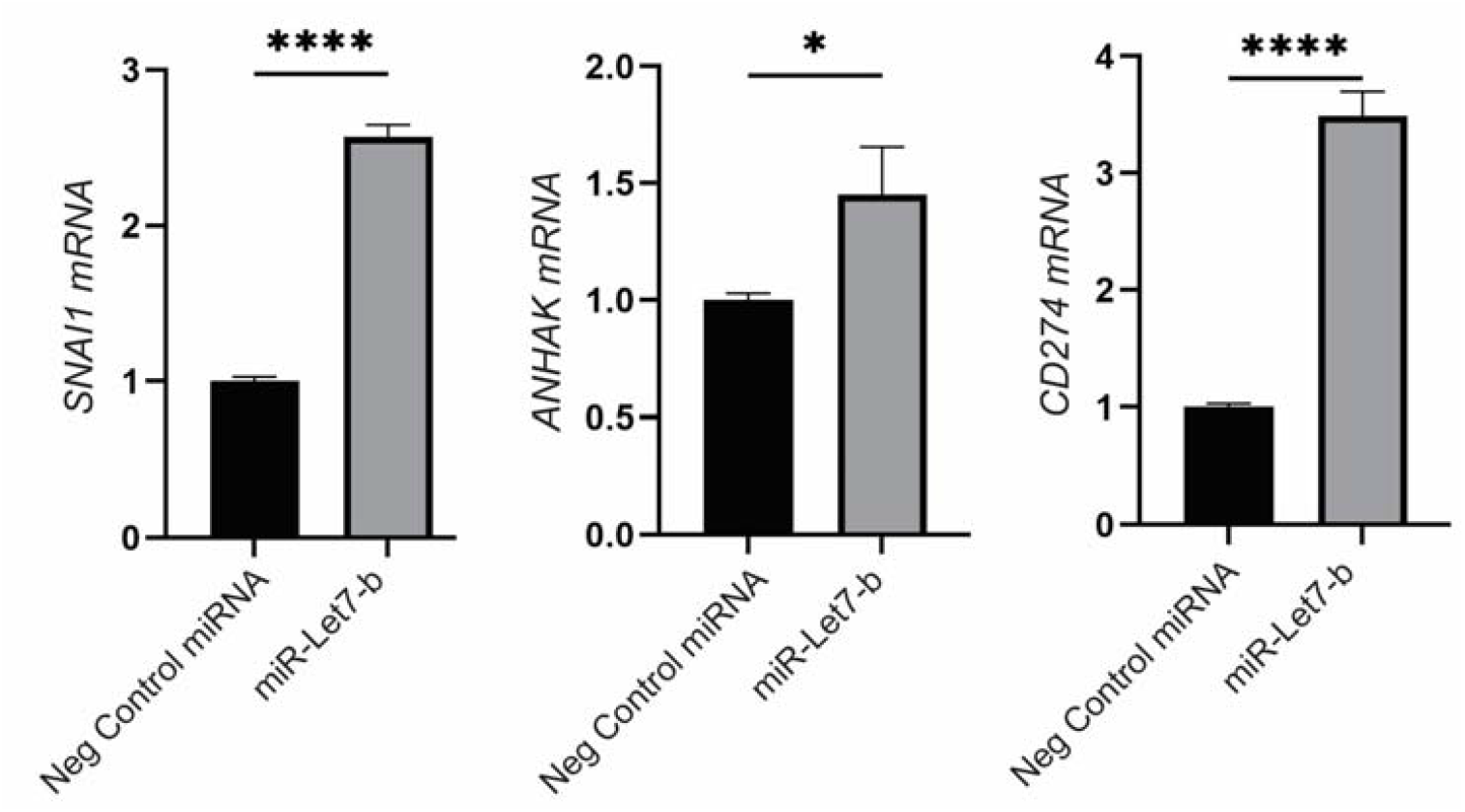
miR-let-7b upregulates selected EMT genes and PD-L1 (*CD274*) in DU145 cells. mRNA expression of selected genes encoding *SNAI1, AHNAK*, and *CD274* was measured by RT-PCR relative to RNA encoding β-actin (*ACTB*). Cells were transfected with miR-let-7b (selected based on its functional significance for EMT and immune exhaustion), or negative control miRNA (non-targeting) at 25 nM and incubated for 48 hours. Data are means ± SEM from three independent experiments; *, *P* < 0.05; ***, *P* < 0.001; and ****, *P* <0.0001 by unpaired two-tailed t-test.

**Supplemental Fig. S7.**
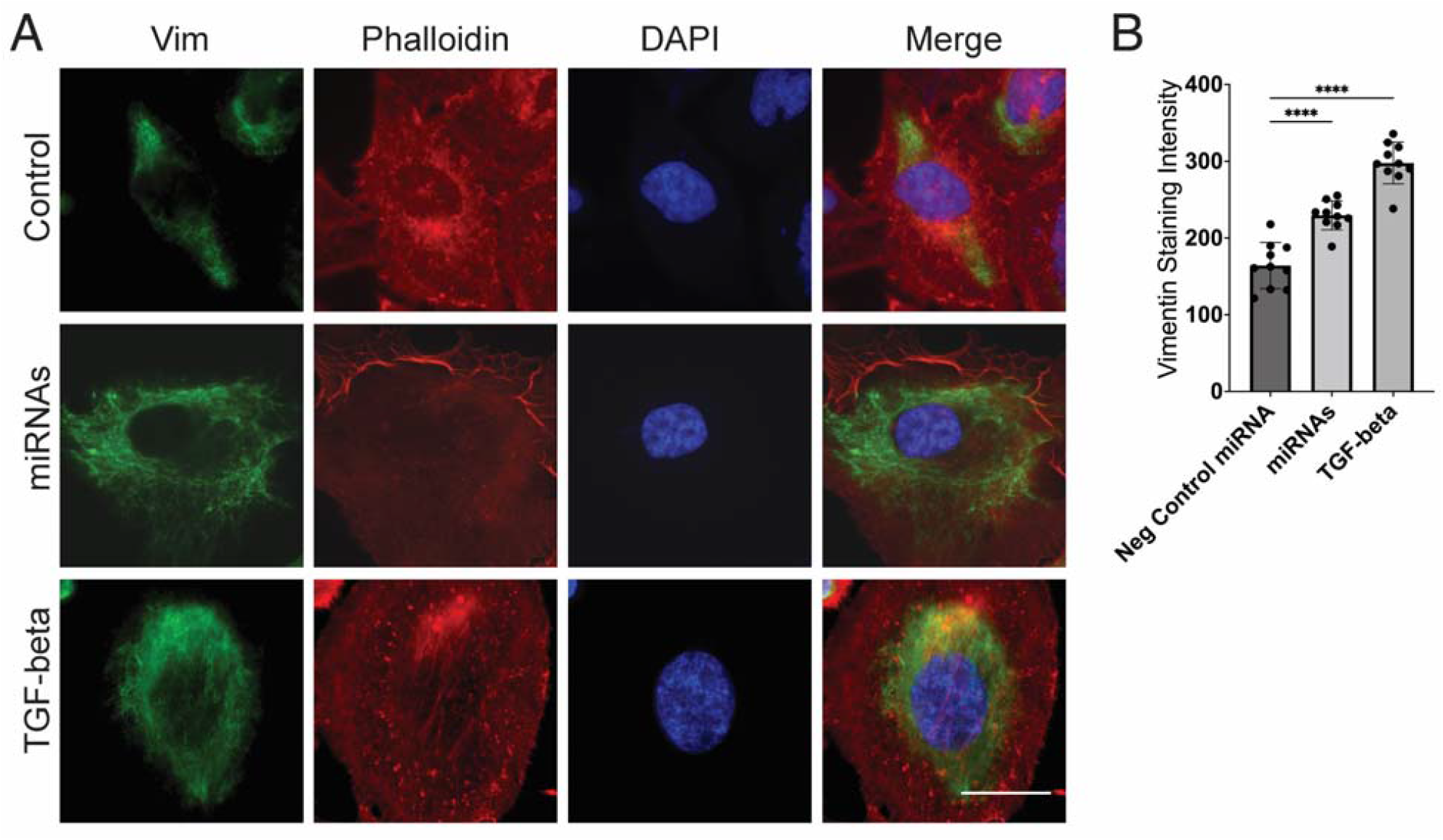
Top three miRNAs upregulate vimentin protein expression in the PC3 model. **(A)** Immunofluorescence staining for vimentin was measured in PC3 cells transfected with an equimolar combination of three miRNAs, miR-let-7b-3p, miR-374-5p and miR-93-5p (each 15 nM, total 45 nM), or negative control miRNA (non-target), at 45 nM, and incubated for 48 hours. Cells were stained with vimentin primary antibody and Alexa Fluor-488 (green) secondary antibody. Cytoskeleton was stained with Alexa Fluor-568 Phalloidin (red) and nucleic DNA was stained with DAPI. Scale bar, 10 µm. For each of N = 3 independent experiments, 10 images were examined by immunofluorescence. One representative image is shown, of 10 images collected for each of the three experimental conditions with three replicates. **(B)** Quantification of vimentin staining intensity in 10 individual cells in each condition. Data analyzed by one-way ANOVA; ****, *P* <0.0001.

**Supplemental Fig. S8.**
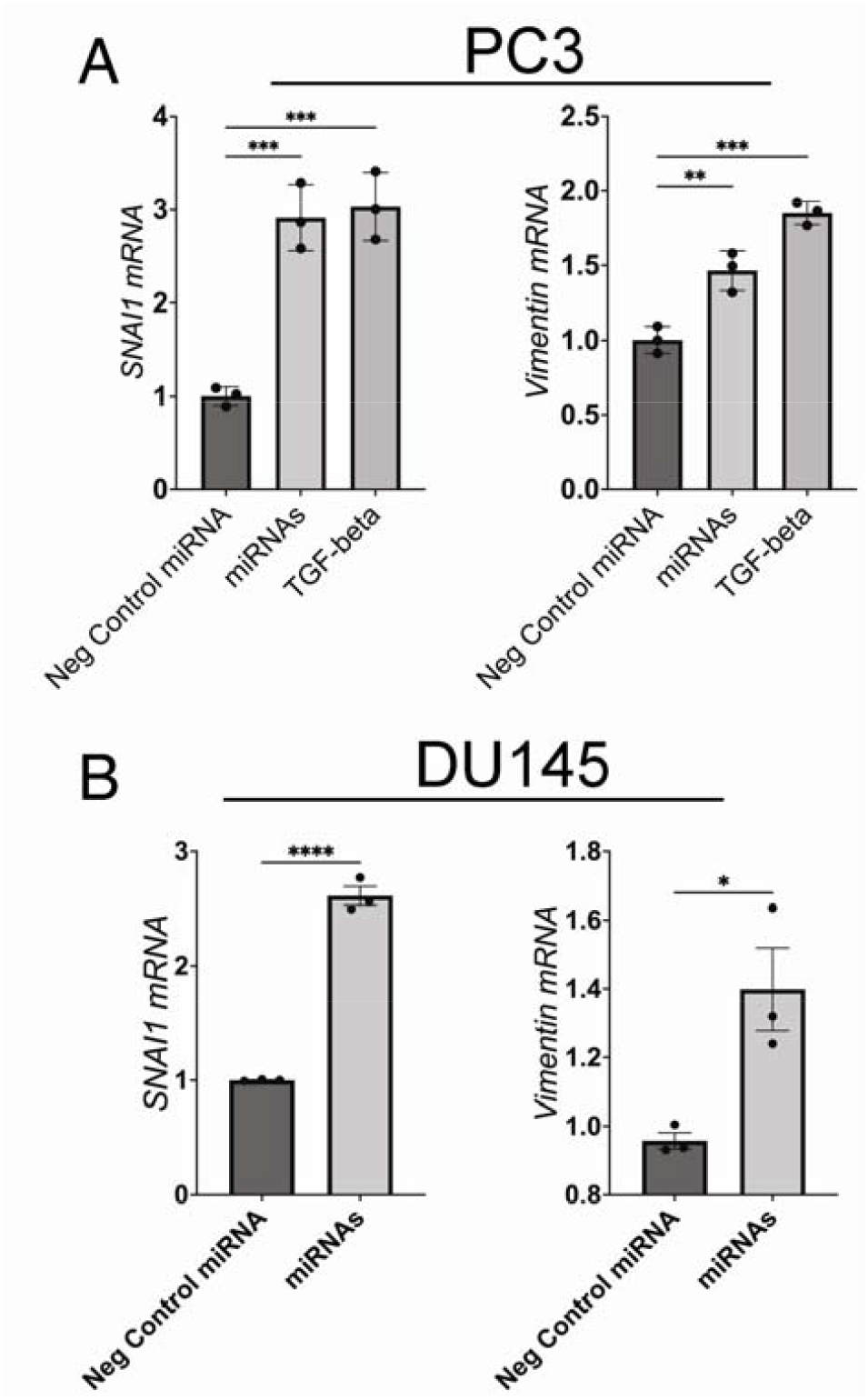
Combination of miR-let-7b-3p, miR-374-5p and miR-93-5p upregulates expression of *SNAI1* and *Vimentin* in human prostate cancer cell lines. PC3 **(A)** and DU145 **(B)** cells were transfected with the equimolar combination of three mRNAs miR-let-7b-3p, miR-374-5p and miR-93-5p (each 15 nM, total 45 nM), or negative control miRNA (non-targeting, 45 nM), and incubated for 48 hours. The mRNA expression of selected genes encoding *SNAI1*, and *Vimentin* was measured by RT-PCR relative to RNA encoding β-actin (ACTB). Data are means ± SEM from three independent experiments; *, *P* < 0.05; **, *P* < 0.01; ***, *P* < 0.001; and ****, *P* <0.0001 by one-way ANOVA in A, and unpaired two-tailed t-test in B. TGFB, TGF-β positive control at 5 ng/mL for *SNAI1* and *Vimentin* induction.

**Supplemental Fig. S9.**
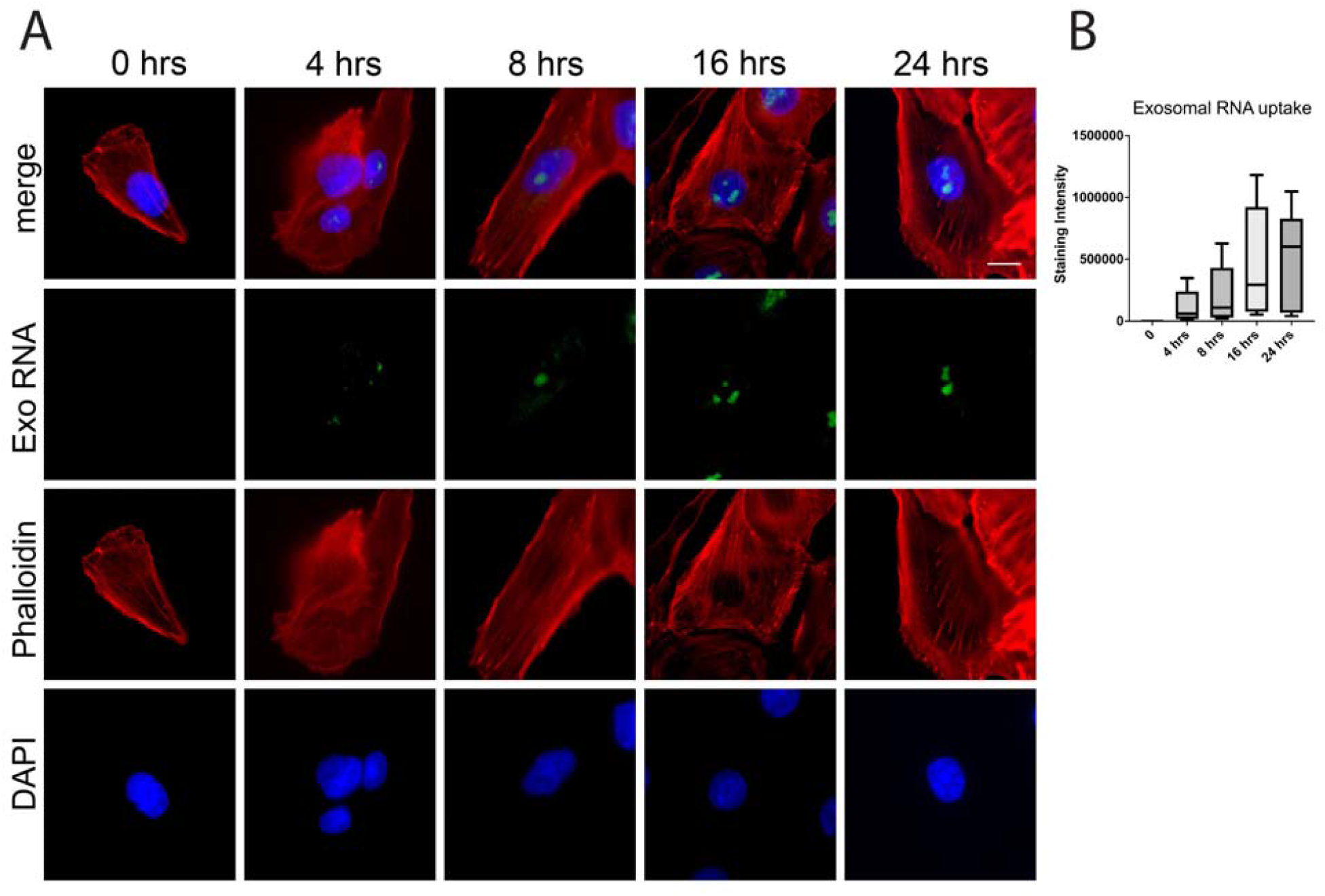
Time course of exosomal RNA uptake into DU145 cell nuclei. (**A**) Exosome RNA was stained with RNASelect (*Exo RNA*; Alexa Fluor™-488; *green*), then exosomes were washed and added to DU145 cells. Cells were imaged at 0, 4, 8, 16 and 24-hour time points. Filamentous actin (F-actin) was stained with Alexa Fluor™ Phalloidin probe 568nm (*Phalloidin*) and DNA was visualized by DAPI counterstain (*DAPI*). One representative field of view is shown, out of 25 images collected for each of the three experimental conditions with three replicates. Scale bar, 10 µm. The mean intensity of stained RNA (Alexa Fluor™-488) was quantified using ImageJ (**B**).

**Supplemental Fig. S10.**
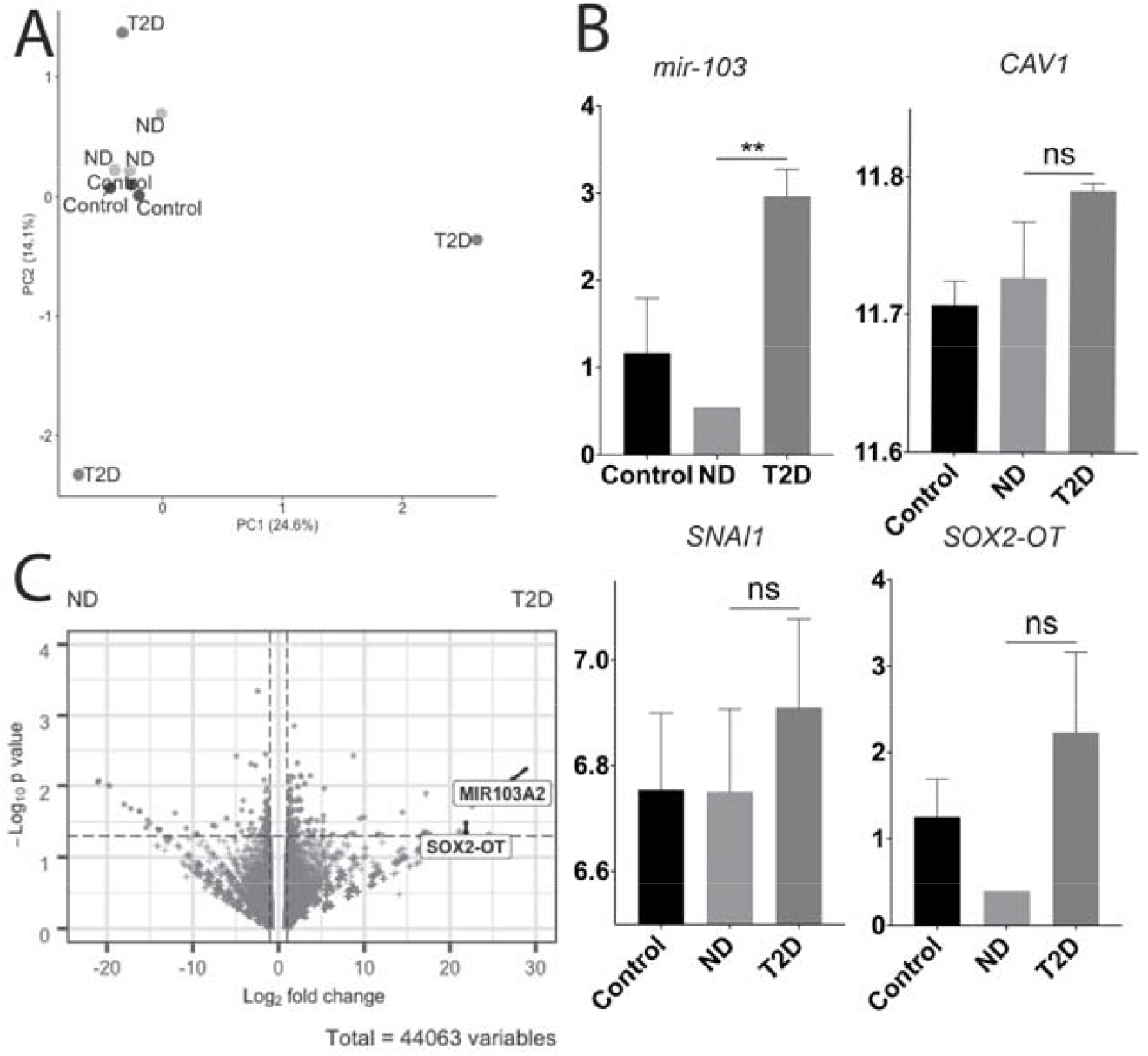
Genome-wide transcriptional analysis by RNA seq. (**A)** Plot of PC2 vs. PC1, computed across all genes in the DU145 samples. Plasma exosomes of T2D subjects had unique and different PCA values, while plasma exosomes of ND subjects had PCA values that were close to the media-only control values. (**B)** RNA seq expression of miR103a, SOX2-OT, Cav1 and SNAI1 in DU145 cells treated with ND and T2D exosomes. The variance stabilizing transformed (VST) expression values for each gene were z-score-normalized to a mean of zero and standard deviation of 1 within all replicates of all samples. The Y axis shows the VST values. (**C)** Differential expression of genes unregulated in DU145 cells treated with T2D exosomes (*right*) vs. those of ND exosomes (*left*). Significantly differentially expressed genes were identified using a *p* value cutoff of 0.05 and a fold change cutoff of 1 (dotted lines). Figure was generated using EnhancedVolcano package.

**Supplemental Fig. S11.**
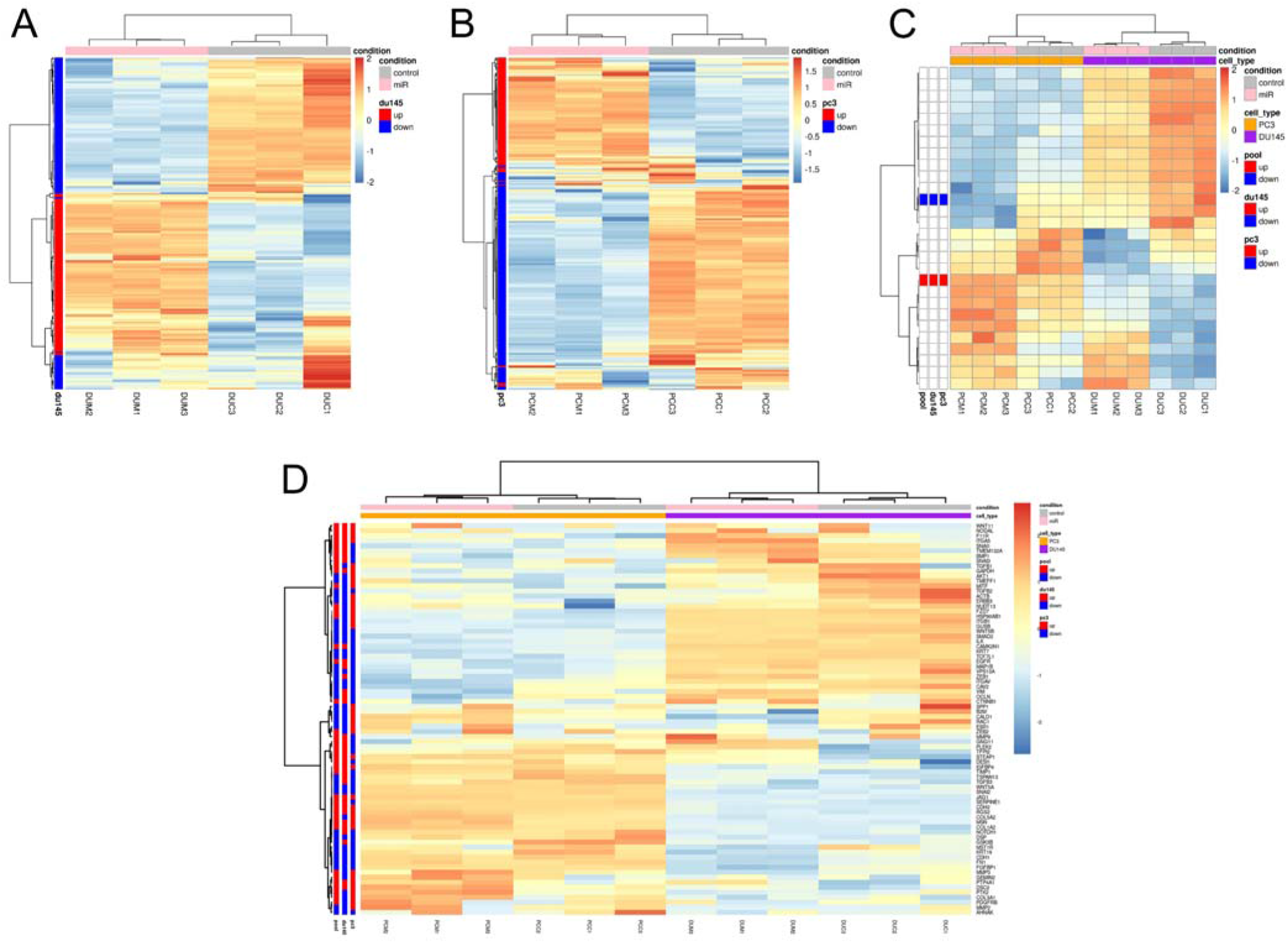
Transcriptional analysis by RNA seq demonstrates enrichment of EMT signature in DU145 and PC3 cell lines, induced by top three T2D exosomal miRNAs. RNA seq analysis of prostate cancer cell lines transfected with the top three miRNA hits (miR-let-7b-3p, miR-374-5p and miR-93-5p) confirmed reproducibility across biological replicates for 439 genes in **(A)** DU145 cells, **(B)** PC3 cells and **(C)** the intersection of the genesets from A and B. Heatmap of upregulated and downregulated genes showed consistency within group and within replicate. **(D)** Enrichment of the EMT hallmark signature from the Fig. 1 Qiagen geneset was confirmed. (DUM1-3, top hit miRNAs in DU145 cells; DUC1-3; non-targeted control miRNAs in DU145 cells; PCM1-3, top hit miRNAs in PC3 cells; PCC1-3, non-targeted control miRNAs in PC3 cells)

## Table legends

**Supplementary Table 1: Proteomics profile of plasma exosomes from 3 ND and 3 T2D subjects**.

Proteins are ranked based on the PSM difference between ND and T2D. (ND, non-diabetic; T2D, Type 2 diabetic; PSMs (number of Peptide-Spectrum Matches), total number of occurrences of unique peptides for each protein; AAs, the number of amino acids in the protein; MW, molecular weight of the protein in kDa; Calc. pI, isoelectric point of each protein)

**Supplementary Table 2: Fold change of T2D vs ND by RNA seq**.

DU145 cells were treated separately with exosome preparations obtained from peripheral blood plasma of three T2D and three ND subjects, as described in Methods, with numbers of exosomes normalized to 10^9^ per well in all cases. After treatment, total RNA of the DU145 cells was isolated and genome-wide transcriptional analysis was conducted by RNA seq. Variance-stabilizing transformation (VST) was accomplished using the “Variance Stabilizing Transformation” function in the DESeq2 R package (version 1.23.10). Significantly differentially expressed genes were identified using a p value cutoff of 0.05 and a fold change cutoff of 1.

## Author Contributions

Conception and design: G.V.D. Methodology: N.J., A.C., M.K., S.M.

Acquisition of data: N.J., M.K, I.R.P., Y.Q., R.Y., P.L., C.E.S.

Analysis and interpretation of data: N.J., A.C., M.K, I.R.P., Y.Q., C.E.S.

Writing, editing, and revision of manuscript: N.J., C.E.S., J.M., K.M., M.H., G.A.G., C.M.H, S.M., G.V.D.

Study supervision: G.V.D.

## Acknowledgements

We thank the Boston University-Boston Medical Center Flow Cytometry Core Facility, the Microarray and Sequencing Resource, and Cellular Imaging Core Facilities for technical assistance. The results shown in this report are based in part upon data generated by the TCGA Research Network: https://www.cancer.gov/tcga.

## References

[1] C. Steele, C. Thomas, henley SJ, Massetti GM, Galuska DA, Agurs-Collins T, Puckett M, Richardson LC, Vital signs: trends in incidence of cancers associated with overweight and obesity-United States, 2014 (2005) 1052–1058.

[2] C.f.D. Control, Prevention, National diabetes statistics report, 2020, Atlanta, GA: Centers for Disease Control and Prevention, US Department of Health and Human Services, (2020) 12–15.

[3] S. Yaturu, Obesity and type 2 diabetes, Journal of diabetes mellitus, 1 (2011) 79–95.

[4] S.T. Fleming, A. Rastogi, A. Dmitrienko, K.D. Johnson, A comprehensive prognostic index to predict survival based on multiple comorbidities: a focus on breast cancer, Medical care, (1999) 601–614.

[5] K.d.C.B. Ribeiro, L.P. Kowalski, M.d.R.D. De Oliveira, Perioperative complications, comorbidities, and survival in oral or oropharyngeal cancer, Archives of Otolaryngology–Head & Neck Surgery, 129 (2003) 219–228.

[6] R. Rieker, E. Hammer, R. Eisele, E. Schmid, J. Högel, The impact of comorbidity on the overall survival and the cause of death in patients after colorectal cancer resection, Langenbeck’s Archives of Surgery, 387 (2002) 72–76.

[7] C. Kastner, J. Armitage, A. Kimble, J. Rawal, P. Carter, S. Venn, The Charlson comorbidity score: a superior comorbidity assessment tool for the prostate cancer multidisciplinary meeting, Prostate cancer and prostatic diseases, 9 (2006) 270–274.

[8] A. Berglund, H. Garmo, C. Tishelman, L. Holmberg, P. Stattin, M. Lambe, Comorbidity, treatment and mortality: a population based cohort study of prostate cancer in PCBaSe Sweden, The Journal of urology, 185 (2011) 833–840.

[9] M. Extermann, Interaction between comorbidity and cancer, Cancer Control, 14 (2007) 13–22.

[10] J. Coebergh, M. Janssen-Heijnen, P. Post, P. Razenberg, Serious co-morbidity among unselected cancer patients newly diagnosed in the southeastern part of The Netherlands in 1993– 1996, Journal of clinical epidemiology, 52 (1999) 1131–1136.

[11] K.-L.C. Jen, K. Brogan, O.G. Washington, J.M. Flack, N.T. Artinian, Poor nutrient intake and high obese rate in an urban African American population with hypertension, Journal of the American College of Nutrition, 26 (2007) 57–65.

[12] R.B. Cadzow, T.J. Servoss, C.H. Fox, The health status of patients of a student-run free medical clinic in inner-city Buffalo, NY, The Journal of the American Board of Family Medicine, 20 (2007) 572–580.

[13] G.A. Nichols, M. McBurnie, L. Paul, J.E. Potter, S. McCann, K. Mayer, G. Melgar, S. D’Amato, J.E. DeVoe, Peer Reviewed: The High Prevalence of Diabetes in a Large Cohort of Patients Drawn From Safety Net Clinics, Preventing chronic disease, 13 (2016).

[14] H.K. Seligman, E.A. Jacobs, A. Lopez, U. Sarkar, J. Tschann, A. Fernandez, Food insecurity and hypoglycemia among safety net patients with diabetes, Archives of Internal Medicine, 171 (2011) 1204–1206.

[15] S.A. Berkowitz, X. Gao, K.L. Tucker, Food-insecure dietary patterns are associated with poor longitudinal glycemic control in diabetes: results from the Boston Puerto Rican Health study, Diabetes care, 37 (2014) 2587–2592.

[16] N.J. Vickers, Animal communication: when i’m calling you, will you answer too?, Current biology, 27 (2017) R713–R715.

[17] C. Muller, Tumour-surrounding adipocytes are active players in breast cancer progression, Annales d’endocrinologie, Elsevier, 2013, pp. 108–110.

[18] L. Lore, H. An, L. Evelyne, V.B. Mieke, V. Jo, M. Dawn, B. Geert, V.D.B. Rudy, M. Cathérine, B. Marc, Secretome analysis of breast cancer-associated adipose tissue to identify paracrine regulators of breast cancer growth, Oncotarget, 8 (2017) 47239.

[19] C. Vaysse, J. Lømo, Ø. Garred, F. Fjeldheim, T. Lofteroed, E. Schlichting, A. McTiernan, H. Frydenberg, A. Husøy, S. Lundgren, Inflammation of mammary adipose tissue occurs in overweight and obese patients exhibiting early-stage breast cancer, NPJ breast cancer, 3 (2017) 1–10.

[20] V. Laurent, A. Guérard, C. Mazerolles, S. Le Gonidec, A. Toulet, L. Nieto, F. Zaidi, B. Majed, D. Garandeau, Y. Socrier, Periprostatic adipocytes act as a driving force for prostate cancer progression in obesity, Nature communications, 7 (2016) 1–15.

[21] V. Laurent, A. Toulet, C. Attané, D. Milhas, S. Dauvillier, F. Zaidi, E. Clement, M. Cinato, S. Le Gonidec, A. Guérard, Periprostatic adipose tissue favors prostate cancer cell invasion in an obesity-dependent manner: role of oxidative stress, Molecular Cancer Research, 17 (2019) 821–835.

[22] D. Estève, M. Roumiguié, C. Manceau, D. Milhas, C. Muller, Periprostatic adipose tissue: A heavy player in prostate cancer progression, Current Opinion in Endocrine and Metabolic Research, 10 (2020) 29–35.

[23] J. Hammarsten, B. Högstedt, Hyperinsulinaemia: a prospective risk factor for lethal clinical prostate cancer, European journal of cancer, 41 (2005) 2887–2895.

[24] J. Ma, H. Li, E. Giovannucci, L. Mucci, W. Qiu, P.L. Nguyen, J.M. Gaziano, M. Pollak, M.J. Stampfer, Prediagnostic body-mass index, plasma C-peptide concentration, and prostate cancer-specific mortality in men with prostate cancer: a long-term survival analysis, The lancet oncology, 9 (2008) 1039–1047.

[25] J.L. Wright, S.R. Plymate, M.P. Porter, J.L. Gore, D.W. Lin, E. Hu, S.B. Zeliadt, Hyperglycemia and prostate cancer recurrence in men treated for localized prostate cancer, Prostate cancer and prostatic diseases, 16 (2013) 204–208.

[26] T. Karantanos, S. Karanika, G. Gignac, Uncontrolled diabetes predicts poor response to novel antiandrogens, Endocrine-related cancer, 23 (2016) 691–698.

[27] L. Magura, R. Blanchard, B. Hope, J.R. Beal, G.G. Schwartz, A.E. Sahmoun, Hypercholesterolemia and prostate cancer: a hospital-based case–control study, Cancer Causes & Control, 19 (2008) 1259–1266.

[28] E. Kheterpal, J.D. Sammon, M. Diaz, A. Bhandari, Q.-D. Trinh, N. Pokala, P. Sharma, M. Menon, P.K. Agarwal, Effect of metabolic syndrome on pathologic features of prostate cancer, Urologic Oncology: Seminars and Original Investigations, Elsevier, 2013, pp. 1054–1059.

[29] J. Flanagan, P.K. Gray, N. Hahn, J. Hayes, L. Myers, C. Carney-Doebbeling, C. Sweeney, Presence of the metabolic syndrome is associated with shorter time to castration-resistant prostate cancer, Annals of Oncology, 22 (2011) 801–807.

[30] I.-C. Yu, H.-Y. Lin, J.D. Sparks, S. Yeh, C. Chang, Androgen receptor roles in insulin resistance and obesity in males: the linkage of androgen-deprivation therapy to metabolic syndrome, Diabetes, 63 (2014) 3180–3188.

[31] A.W. Wyatt, M. Annala, R. Aggarwal, K. Beja, F. Feng, J. Youngren, A. Foye, P. Lloyd, M. Nykter, T.M. Beer, Concordance of circulating tumor DNA and matched metastatic tissue biopsy in prostate cancer, JNCI: Journal of the National Cancer Institute, 109 (2017).

[32] R. Kanwal, A.R. Plaga, X. Liu, G.C. Shukla, S. Gupta, MicroRNAs in prostate cancer: Functional role as biomarkers, Cancer Letters, 407 (2017) 9–20.

[33] A.M. Aghdam, A. Amiri, R. Salarinia, A. Masoudifar, F. Ghasemi, H. Mirzaei, MicroRNAs as diagnostic, prognostic, and therapeutic biomarkers in prostate cancer, Critical Reviews™ in Eukaryotic Gene Expression, 29 (2019).

[34] A. Watahiki, R.J. Macfarlane, M.E. Gleave, F. Crea, Y. Wang, C.D. Helgason, K.N. Chi, Plasma miRNAs as biomarkers to identify patients with castration-resistant metastatic prostate cancer, International journal of molecular sciences, 14 (2013) 7757–7770.

[35] G. Rabinowits, C. Gerçel-Taylor, J.M. Day, D.D. Taylor, G.H. Kloecker, Exosomal microRNA: a diagnostic marker for lung cancer, Clinical lung cancer, 10 (2009) 42–46.

[36] A. Michael, S.D. Bajracharya, P.S. Yuen, H. Zhou, R.A. Star, G.G. Illei, I. Alevizos, Exosomes from human saliva as a source of microRNA biomarkers, Oral diseases, 16 (2010) 34–38.

[37] J. Skog, T. Würdinger, S. van Rijn, D. Meijer, L. Gainche, Glioblastoma microvesicles transport RNA and proteins that promote tumour growth and provide diagnostic biomarkers. NatCellBiol10 (12): 1470–6.[10.1038/ncb1800] Chapter 1 of tumor-derived exosomes as diagnostic biomarkers of ovarian cancer, Gynecol Oncol, 110 (2008) 13–21.

[38] R. Nedaeinia, M. Manian, M. Jazayeri, M. Ranjbar, R. Salehi, M. Sharifi, F. Mohaghegh, M. Goli, S. Jahednia, A. Avan, Circulating exosomes and exosomal microRNAs as biomarkers in gastrointestinal cancer, Cancer gene therapy, 24 (2017) 48–56.

[39] N. Kosaka, H. Iguchi, T. Ochiya, Circulating microRNA in body fluid: a new potential biomarker for cancer diagnosis and prognosis, Cancer science, 101 (2010) 2087–2092.

[40] T. Matsumura, K. Sugimachi, H. Iinuma, Y. Takahashi, J. Kurashige, G. Sawada, M. Ueda, R. Uchi, H. Ueo, Y. Takano, Exosomal microRNA in serum is a novel biomarker of recurrence in human colorectal cancer, British journal of cancer, 113 (2015) 275–281.

[41] S.J. Freedland, W.J. Aronson, Examining the relationship between obesity and prostate cancer, Reviews in urology, 6 (2004) 73.

[42] K. Di Sebastiano, J. Pinthus, W. Duivenvoorden, M. Mourtzakis, Glucose impairments and insulin resistance in prostate cancer: The role of obesity, nutrition and exercise, Obesity Reviews, 19 (2018) 1008–1016.

[43] F. Nik-Ahd, L.E. Howard, A.T. Eisenberg, W.J. Aronson, M.K. Terris, M.R. Cooperberg, C.L. Amling, C.J. Kane, S.J. Freedland, Poorly controlled diabetes increases the risk of metastases and castration-resistant prostate cancer in men undergoing radical prostatectomy: Results from the SEARCH database, Cancer, 125 (2019) 2861–2867.

[44] S. Kelkar, T. Oyekunle, A. Eisenberg, L. Howard, W.J. Aronson, C.J. Kane, C.L. Amling, M.R. Cooperberg, Z. Klaassen, M.K. Terris, Diabetes and Prostate Cancer Outcomes in Obese and Nonobese Men After Radical Prostatectomy, JNCI Cancer Spectrum, 5 (2021) pkab023.

[45] M.F. Sona, S.-K. Myung, K. Park, G. Jargalsaikhan, Type 1 diabetes mellitus and risk of cancer: a meta-analysis of observational studies, Japanese journal of clinical oncology, 48 (2018) 426–433.

[46] R. Peila, T.E. Rohan, Diabetes, glycated hemoglobin, and risk of cancer in the UK biobank study, Cancer Epidemiology and Prevention Biomarkers, 29 (2020) 1107–1119.

[47] A.J. Burton, K.M. Tilling, J.M. Holly, F.C. Hamdy, M.-A.E. Rowlands, J.L. Donovan, R.M. Martin, Metabolic imbalance and prostate cancer progression, International journal of molecular epidemiology and genetics, 1 (2010) 248.

[48] A.J. Klil-Drori, L. Azoulay, M.N. Pollak, Cancer, obesity, diabetes, and antidiabetic drugs: is the fog clearing?, Nature reviews Clinical oncology, 14 (2017) 85–99.

[49] N. Jafari, M. Kolla, T. Meshulam, J.S. Shafran, Y. Qiu, A.N. Casey, I.R. Pompa, C.S. Ennis, C.S. Mazzeo, N. Rabhi, S.R. Farmer, G.V. Denis, Adipocyte-derived exosomes may promote breast cancer progression in type 2 diabetes, Science Signaling, 14 (2021) eabj2807.

[50] B. Zhang, Y. Yang, X. Shi, W. Liao, M. Chen, A.S.-L. Cheng, H. Yan, C. Fang, S. Zhang, G. Xu, Proton pump inhibitor pantoprazole abrogates adriamycin-resistant gastric cancer cell invasiveness via suppression of Akt/GSK-β/β-catenin signaling and epithelial–mesenchymal transition, Cancer letters, 356 (2015) 704–712.

[51] J.S. Shafran, G.P. Andrieu, B. Györffy, G.V. Denis, BRD4 regulates metastatic potential of castration-resistant prostate cancer through AHNAK, Molecular Cancer Research, 17 (2019) 1627–1638.

[52] J.S. Shafran, N. Jafari, A.N. Casey, B. Győrffy, G.V. Denis, BRD4 regulates key transcription factors that drive epithelial–mesenchymal transition in castration-resistant prostate cancer, Prostate cancer and prostatic diseases, 24 (2021) 268–277.

[53] G.P. Andrieu, J.S. Shafran, C.L. Smith, A.C. Belkina, A.N. Casey, N. Jafari, G.V. Denis, BET protein targeting suppresses the PD-1/PD-L1 pathway in triple-negative breast cancer and elicits anti-tumor immune response, Cancer letters, 465 (2019) 45–58.

[54] A. Dobin, C.A. Davis, F. Schlesinger, J. Drenkow, C. Zaleski, S. Jha, P. Batut, M. Chaisson, T.R. Gingeras, STAR: ultrafast universal RNA-seq aligner, Bioinformatics, 29 (2013) 15–21.

[55] M.I. Love, W. Huber, S. Anders, Moderated estimation of fold change and dispersion for RNA-seq data with DESeq2, Genome biology, 15 (2014) 1–21.

[56] Y. Lou, L. Diao, E.R.P. Cuentas, W.L. Denning, L. Chen, Y.H. Fan, L.A. Byers, J. Wang, V.A. Papadimitrakopoulou, C. Behrens, Epithelial–mesenchymal transition is associated with a distinct tumor microenvironment including elevation of inflammatory signals and multiple immune checkpoints in lung adenocarcinoma, Clinical Cancer Research, 22 (2016) 3630–3642.

[57] J.C. Thompson, W.-T. Hwang, C. Davis, C. Deshpande, S. Jeffries, Y. Rajpurohit, V. Krishna, D. Smirnov, R. Verona, M.V. Lorenzi, Gene signatures of tumor inflammation and epithelial-to-mesenchymal transition (EMT) predict responses to immune checkpoint blockade in lung cancer with high accuracy, Lung Cancer, 139 (2020) 1–8.

[58] M. Taki, K. Abiko, M. Ukita, R. Murakami, K. Yamanoi, K. Yamaguchi, J. Hamanishi, T. Baba, N. Matsumura, M. Mandai, Tumor Immune Microenvironment during Epithelial– Mesenchymal Transition, Clinical Cancer Research, 27 (2021) 4669–4679.

[59] Q. Lin, C.-R. Zhou, M.-J. Bai, D. Zhu, J.-W. Chen, H.-F. Wang, M.-A. Li, C. Wu, Z.-R. Li, M.-S. Huang, Exosome-mediated miRNA delivery promotes liver cancer EMT and metastasis, American journal of translational research, 12 (2020) 1080.

[60] D. Son, Y. Kim, S. Lim, H.-G. Kang, D.-H. Kim, J.W. Park, W. Cheong, H.K. Kong, W. Han, W.-Y. Park, miR-374a-5p promotes tumor progression by targeting ARRB1 in triple negative breast cancer, Cancer letters, 454 (2019) 224–233.

[61] L. Liang, L. Zhao, Y. Zan, Q. Zhu, J. Ren, X. Zhao, MiR-93-5p enhances growth and angiogenesis capacity of HUVECs by down-regulating EPLIN, Oncotarget, 8 (2017) 107033.

[62] Y. Yang, B. Jia, X. Zhao, Y. Wang, W. Ye, miR-93-5p may be an important oncogene in prostate cancer by bioinformatics analysis, Journal of cellular biochemistry, 120 (2019) 10463–10483.

[63] S. Rana, G.N. Valbuena, E. Curry, C.L. Bevan, H.C. Keun, MicroRNAs as biomarkers for prostate cancer prognosis: a systematic review and a systematic reanalysis of public data, British journal of cancer, (2022) 1–12.

[64] Z. Wang, L. Xu, Y. Hu, Y. Huang, Y. Zhang, X. Zheng, S. Wang, Y. Wang, Y. Yu, M. Zhang, miRNA let-7b modulates macrophage polarization and enhances tumor-associated macrophages to promote angiogenesis and mobility in prostate cancer, Scientific reports, 6 (2016) 1–11.

[65] X. Liang, Z. Li, Q. Men, Y. Li, H. Li, T. Chong, miR-326 functions as a tumor suppressor in human prostatic carcinoma by targeting Mucin1, Biomedicine & Pharmacotherapy, 108 (2018) 574–583.

[66] K. Kang, J. Zhang, X. Zhang, Z. Chen, MicroRNA-326 inhibits melanoma progression by targeting KRAS and suppressing the AKT and ERK signalling pathways, Oncology reports, 39 (2018) 401–410.

[67] G. Andrieu, A.H. Tran, K.J. Strissel, G.V. Denis, BRD4 regulates breast cancer dissemination through Jagged1/Notch1 signaling, Cancer research, 76 (2016) 6555–6567.

[68] G.P. Andrieu, G.V. Denis, BET proteins exhibit transcriptional and functional opposition in the epithelial-to-mesenchymal transition, Molecular Cancer Research, 16 (2018) 580–586.

[69] S.J. Hogg, S.J. Vervoort, S. Deswal, C.J. Ott, J. Li, L.A. Cluse, P.A. Beavis, P.K. Darcy, B.P. Martin, A. Spencer, BET-bromodomain inhibitors engage the host immune system and regulate expression of the immune checkpoint ligand PD-L1, Cell reports, 18 (2017) 2162–2174.

[70] X. Wang, Y. Zhou, Y. Peng, T. Huang, F. Xia, T. Yang, Q. Duan, W. Zhang, Bromodomain-containing protein 4 contributes to renal fibrosis through the induction of epithelial-mesenchymal transition, Experimental cell research, 383 (2019) 111507.

[71] X. Jing, S. Shao, Y. Zhang, A. Luo, L. Zhao, L. Zhang, S. Gu, X. Zhao, BRD4 inhibition suppresses PD-L1 expression in triple-negative breast cancer, Experimental cell research, 392 (2020) 112034.

[72] M.C. Haffner, W. Zwart, M.P. Roudier, L.D. True, W.G. Nelson, J.I. Epstein, A.M. De Marzo, P.S. Nelson, S. Yegnasubramanian, Genomic and phenotypic heterogeneity in prostate cancer, Nature Reviews Urology, 18 (2021) 79–92.

[73] O. Sartor, J.S. de Bono, Metastatic prostate cancer, New England Journal of Medicine, 378 (2018) 645–657.

[74] W.J. Catalona, D.S. Smith, T.L. Ratliff, K.M. Dodds, D.E. Coplen, J.J. Yuan, J.A. Petros, G.L. Andriole, Measurement of prostate-specific antigen in serum as a screening test for prostate cancer, New England Journal of Medicine, 324 (1991) 1156–1161.

[75] R. Etzioni, A. Tsodikov, A. Mariotto, A. Szabo, S. Falcon, J. Wegelin, D. Ditommaso, K. Karnofski, R. Gulati, D.F. Penson, Quantifying the role of PSA screening in the US prostate cancer mortality decline, Cancer Causes & Control, 19 (2008) 175–181.

[76] B. Djavan, A. Zlotta, M. Remzi, K. Ghawidel, A. Basharkhah, C.C. Schulman, M. Marberger, Optimal predictors of prostate cancer on repeat prostate biopsy: a prospective study of 1,051 men, The Journal of urology, 163 (2000) 1144–1149.

[77] I.M. Thompson, D.K. Pauler, P.J. Goodman, C.M. Tangen, M.S. Lucia, H.L. Parnes, L.M. Minasian, L.G. Ford, S.M. Lippman, E.D. Crawford, Prevalence of prostate cancer among men with a prostate-specific antigen level≤ 4.0 ng per milliliter, New England Journal of Medicine, 350 (2004) 2239–2246.

[78] H.Y. Chen, Y.M. Lin, H.C. Chung, Y.D. Lang, C.J. Lin, J. Huang, W.C. Wang, F.M. Lin, Z. Chen, H.D. Huang, miR-103/107 promote metastasis of colorectal cancer by targeting the metastasis suppressors DAPK and KLF4, Cancer research, 72 (2012) 3631–3641.

[79] D. Gao, B. Hou, D. Zhou, Q. Liu, K. Zhang, X. Lu, J. Zhang, H. Zheng, J. Dai, Tumor-derived exosomal miR-103a-2-5p facilitates esophageal squamous cell carcinoma cell proliferation and migration, Eur. Rev. Med. Pharmacol. Sci, 24 (2020) 6097–6110.

[80] X. Liu, Y. Cao, Y. Zhang, H. Zhou, H. Li, Regulatory effect of MiR103 on proliferation, EMT and invasion of oral squamous carcinoma cell through SALL4, Eur. Rev. Med. Pharmacol. Sci, 23 (2019) 9931–9938.

[81] Q. Wo, D. Zhang, L. Hu, J. Lyu, F. Xiang, W. Zheng, J. Shou, X. Qi, Long noncoding RNA SOX2-OT facilitates prostate cancer cell proliferation and migration via miR-369-3p/CFL2 axis, Biochemical and biophysical research communications, 520 (2019) 586–593.

[82] X. Song, H. Wang, J. Wu, Y. Sun, Long Noncoding RNA SOX2-OT Knockdown Inhibits Proliferation and Metastasis of Prostate Cancer Cells Through Modulating the miR-452-5p/HMGB3 Axis and Inactivating Wnt/β-Catenin Pathway, Cancer biotherapy & radiopharmaceuticals, 35 (2020) 682–695.

[83] E.D. Kwon, C.G. Drake, H.I. Scher, K. Fizazi, A. Bossi, A.J. Van den Eertwegh, M. Krainer, N. Houede, R. Santos, H. Mahammedi, Ipilimumab versus placebo after radiotherapy in patients with metastatic castration-resistant prostate cancer that had progressed after docetaxel chemotherapy (CA184-043): a multicentre, randomised, double-blind, phase 3 trial, The lancet oncology, 15 (2014) 700–712.

[84] T. Powles, K.C. Yuen, S. Gillessen, E.E. Kadel, D. Rathkopf, N. Matsubara, C.G. Drake, K. Fizazi, J.M. Piulats, P.J. Wysocki, Atezolizumab with enzalutamide versus enzalutamide alone in metastatic castration-resistant prostate cancer: a randomized phase 3 trial, Nature medicine, (2022) 1–10.

[85] Hänzelmann, S., Castelo, R. & Guinney, J. GSVA: gene set variation analysis for microarray and RNA-Seq data. BMC Bioinformatics 14, 7 (2013).

